# EMBER: A Genome-Scale Approach for a Systematic Characterization of Bacterial Metabolic Heterogeneity through the Growth–Adaptation Trade-Off

**DOI:** 10.64898/2025.12.10.693180

**Authors:** Alvaro Garantilla Becerra, Juan Nogales

**Affiliations:** Systems Biotechnology Group, Department of Systems Biology, Centro Nacional de Biotecnología, CSIC, Madrid, Spain; CNB DNA Biofoundry (CNBio), CSIC, Madrid, Spain; Interdisciplinary Platform for Sustainable Plastics towards a Circular Economy-Spanish National Research Council (SusPlast-CSIC), Madrid, Spain

## Abstract

Microorganisms maintain resilience in fluctuating environments by operating close to a multi-optimality state, balancing growth rate and adaptability. This trade-off dictates bacterial resilience and often complicates metabolic engineering efforts. Addressing it requires identifying specific pathways responsible for diverting metabolic resources away from desired production goals. For this purpose, we introduce EMBER (Exploration of Metabolic trade-offs Based on the mapping of Expression patterns to Reactions), a novel Genome-scale Metabolic Model (GEM) contextualization approach. EMBER integrates transcriptomic data and flux analysis to computationally distinguish between growth-associated reactions (BARs) and adaptive, non-biomass reactions (NBRs). We applied this framework to analyze the metabolic architectures of three diverse and biotechnologically relevant organisms—P. putida, E. coli, and Synechocystis—across various environmental conditions. We revealed marked variability in adaptive resource allocation, with the heterotrophs dedicating substantially more active genes to NBRs (up to 31%) than the photoautotroph Synechocystis (17.5%). Functional analysis showed that BARs consistently supported core metabolism, while NBRs encoded context-specific adaptive functions aligned with the organism’s native environment. Analysis of NBR gene expression variability further suggested that P. putida relies predominantly on Bet-Hedging strategies, whereas E. coli employs more regulated Responsive Switching mechanisms. Overall, EMBER offers a powerful systems biology tool to quantify and functionally interpret metabolic heterogeneity. This systematic identification of NBRs will facilitate precise metabolic engineering efforts via reducing unnecessary fitness costs or harnessing the population heterogeneity for deploying complex biotechnological tasks.

## 1. Introduction

Bacterial colonies are remarkable for their ability to thrive in fluctuating and often hostile environments. A key to this resilience lies in their metabolic flexibility and the emergence of phenotypic heterogeneity, even within clonal populations^1–3^. This cellular individuality is a crucial modulator of phenotypes at the population level and can significantly influence outcomes of critical importance such as survival against antibiotics^2,4^. By taking advantage of this feature, microorganisms adapt to frequent environmental changes through population diversification to survive both short-term perturbations and permanent changes^5,6^.

Broadly, metabolic heterogeneity can be generated by two general mechanisms. First, the cells could “anticipate” the environmental change by stochastically switching phenotype at any time. This is considered a risk-spreading strategy where isogenic populations stochastically generate phenotypes with different fitness-related traits, often resulting in "maladapted individuals" with lower reproductive success in a specific environment. However, these seemingly less fit individuals can have a selective advantage upon a sudden, unpredictable environmental shift, allowing the population to persist in dynamic habitats, albeit with a fitness cost^6^. In this case, adaptation to a new carbon source would be a passive process accomplished by cells that switched phenotypes prior to the environmental change. This behavior is called Bet-Hedging (BH), which is known to maintain phenotypic diversity within a population to ensure survival under unpredictable conditions. Secondly, an initially homogeneous population could actively respond to environmental change with phenotype diversification, where specific subpopulations of the colony adapt to growth under the new conditions. This is often tied to "responsive switching" (RS), which relies on stress-specific sensory circuits. This mechanism allows for a coordinated response of the bacterial population, known as division of labor, where different cells specialize in different tasks^1,6–8^. However, it can lead to an adaptive lag if the environment fluctuates rapidly, where BH would be more advantageous.

As of today, it is assumed that the emergence of heterogeneity in bacterial populations is driven by both intrinsic and extrinsic factors. Intrinsically, stochasticity in gene expression becomes critical, with several studies pointing out how the stochasticity in gene expression can lead to different phenotypes, contributing to the formation of stable subpopulations^5,6^. Extrinsically, environmental influences such as chemical gradients, spatial constraints, and fluctuating conditions are proposed to be key drivers due to their importance in shaping spatial gradients and enabling local adaptation. Moreover, the persistence of diverse metabolic strategies and phenotypes is observed even within clonal populations, that could be attributed to redundancy in metabolic pathways and stress response mechanisms^1–3,8,9^. However, likely due to the lack of experimental tools, relatively little is known about how the propagation of those molecular noises affects the maintenance of metabolism heterogeneity^5^.

As a result of this knowledge gap, several theoretical frameworks regarding metabolic variability have been proposed. We are going to focus on the one developed by Schuetz et al, regarding the multioptimality of bacterial metabolism^10^. This concept refers to the existence of several metabolic flux distributions or states that are equally optimal under given conditions, differing in their physiological or ecological implications. In the developed study, the authors demonstrated that bacterial metabolism operates near a Pareto-optimal surface defined by competing objectives such as ATP yield, biomass production, and minimal flux adjustment. This multidimensional optimal space implies that no single flux distribution is universally better; instead, cells must navigate trade-offs between condition efficiency and adaptability. In *Escherichia coli*, for example, flux distributions cluster in distinct regions of this Pareto surface depending on environmental conditions. Importantly, wild-type strains tend to operate slightly below the Pareto surface, allowing for greater variability in fluxes while maintaining near-optimal performance^10^.

This variability appears not to be random but rather to be evolutionarily selected to minimize the metabolic cost of switching between conditions—a principle termed minimal flux adjustment. Hence, the concept of minimal flux adjustment may explain why bacteria favor flux distributions that are not strictly optimal nor rigid for a single condition but robust across multiple conditions. This translates into the stochasticity of gene expression being favored across bacterial evolution. Also, it provides a hallmark of adaptation through bet-hedging: rather than optimizing for the present, cells prepare for the future. Thus, assuming the previous theory as a driver of evolutionary strategies that promote survival in fluctuating or unpredictably changing environments (Figure 1), we propose a computational approach to characterize their consequences as adaptative responses.

**Figure 1.**
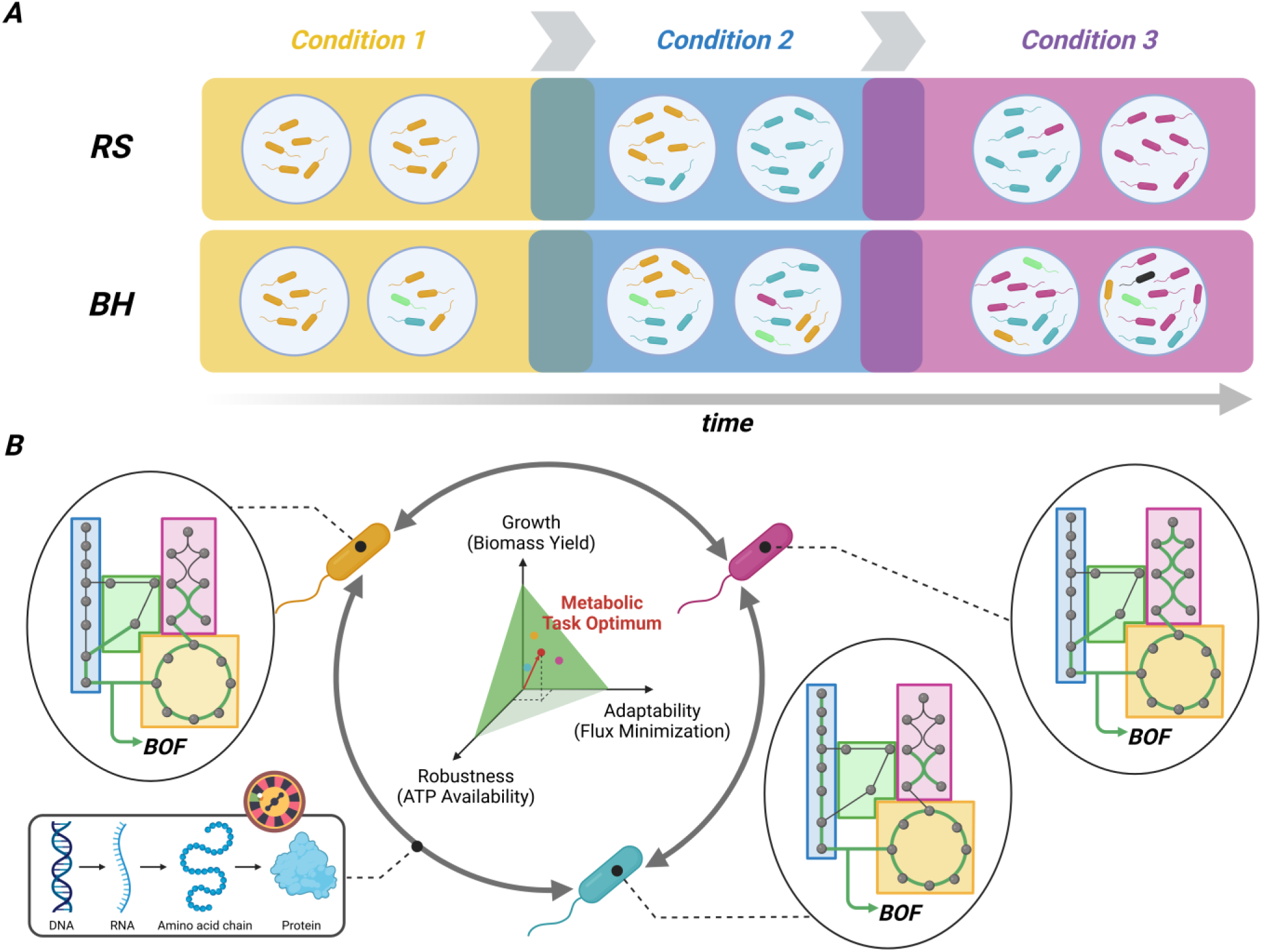
Schematic representation of adaptability-driven heterogeneity of bacterial metabolism. The figure shows an idealized transition between 3 different bacterial metabolic states (colors) according solely to one of the proposed mechanisms, responsive switch or bet-hedging mechanisms (A). According to the multioptimality hypothesis, the metabolic heterogeneity generated either way is stabilized and maintained through several metabolic trade-offs, giving rise to the predominancy of suboptimal, heterogeneous metabolic states (yellow, blue and magenta dots and bacteria) over an optimal phenotype (red dot) for a given metabolic task (B). The network represents the metabolism of the idealized organism, the edges being reactions and the dots representing metabolites. Colored rectangles represent sets of reactions necessary for a metabolic function. Note that, despite the few differences in active reactions (green edges), these changes can imply changes in metabolic and ecological function. This highlights that the noise generated by the stochasticity in gene expression (bottom-left corner) can favor phenotype diversification.

For such a task, we will use genome-scale metabolic models (GEMs) as a mechanistic and computational representation of the bacterial metabolism. GEMs represent a cornerstone in modern systems biology, offering a powerful computational framework to simulate and analyze the intricate metabolic processes within an organism at a systems level. These models provide a comprehensive description of the entire set of biochemical reactions occurring within a cell, integrating vast amounts of genome annotation and experimental data into a mathematically tractable format^11,12^. Since the first genome-scale metabolic reconstruction for *Haemophilus influenzae RD* in 1999, the field has seen an exponential growth in the number of reconstructed models, now encompassing several bacteria, archaea, and eukaryotes (see BiggModel database, http://bigg.ucsd.edu/).

The fundamental purpose of GEMs is to establish a direct link between an organism’s genotype and its observable phenotype, thereby enabling researchers to analyze and contextualize metabolic capabilities with unprecedented detail. They are crucial for understanding how microorganisms adapt and reprogram their metabolic systems in response to various environmental changes^12^. In the broader context of systems biology, GEMs are an indispensable tool for deciphering complex cellular networks and guiding subsequent laboratory work in an iterative manner. They provide a robust scaffold for *in silico* cell metabolism analysis and manipulation, a critical need when traditional experimental approaches alone prove insufficient or too time-consuming for complex biological systems. The successes of these models have tangible implications across diverse fields, including microbial evolution, interaction networks, genetic engineering, and drug discovery^11,13–15^.

## 2. Results

### Assessing the Active Subnetwork of Reactions Underpinning Adaptive Responses

A GEM-based methodology to differentiate between resources allocated to biomass synthesis and those directed toward the implementation of adaptive responses was developed in this work. For this task, it was assumed that the transcriptional response of an organism under a given condition reflects the genetic regulation shaped by evolutionary selection. Post-transcriptional processes, which also contribute to the final adaptive response, were not considered. This decision is justified because transcription already entails a significant resource expenditure, and the primary objective of this study was to identify adaptive responses that divert resources. Consequently, resource allocation in this work was inferred exclusively from transcriptional data. Further developments should also focus on post-transcriptional processes.

Considering this, we focus on GEMs transcriptional contextualization methods that fine-tune GEMs to a given experimental condition according to its transcriptomic response. Specifically, a GIMME-based approach was selected, that excels at determining whether reactions are active or not in a given condition^16^. Basically, GIMME introduces a percentile-based threshold that is mapped through GPRs to infer “*reaction activity”* from gene expression. It also accounts for a given objective that must be maintained (such as BOF). If the objective is not achieved when all reactions determined to be inactive are removed, some of those reactions are added until the objective becomes feasible.

Fueled by GIMME, we could develop EMBER (from <u>E</u>xploration of <u>M</u>etabolic trade-offs <u>B</u>ased on the mapping of <u>E</u>xpression patterns to <u>R</u>eactions) to differentiate between BARs and NBRs genes. EMBER consists of a 2-level contextualization approach (Figure 2). It first performs a direct mapping of genes to reaction expression without considering any GEM objective. Then, it executes a GIMME run, setting the objective so that the biomass objective function (BOF) reaches at least 90% of the wild-type value. These steps yield four distinct sets of reactions: i) Active reactions considering the BOF objective (AG), ii) Active reactions considering only gene expression data (AT), iii) Inactive reactions under the BOF objective (IG), and iv) Inactive reactions considering only expression (IT). From these datasets, reactions classified as inactive by both analyses (IT and IG) were removed from the model. The resulting set of reactions—comprising active reactions (AT and AG) and those classified as inactive by only one of the analyses (IT or IG)—constituted the final contextualized GEM. Subsequently, FVA was applied to identify blocked and non-blocked reactions. Since FVA assumes the biomass objective function (BOF) as the optimization target, reactions classified as blocked under this criterion, even if transcriptionally expressed, were predicted to be unrelated to cell growth. Therefore, these blocked reactions were classified as NBRs, assumed to be active due to adaptation requirements rather than biomass generation. The rest of reactions remaining in the network were classified as BARs.

**Figure 2.**
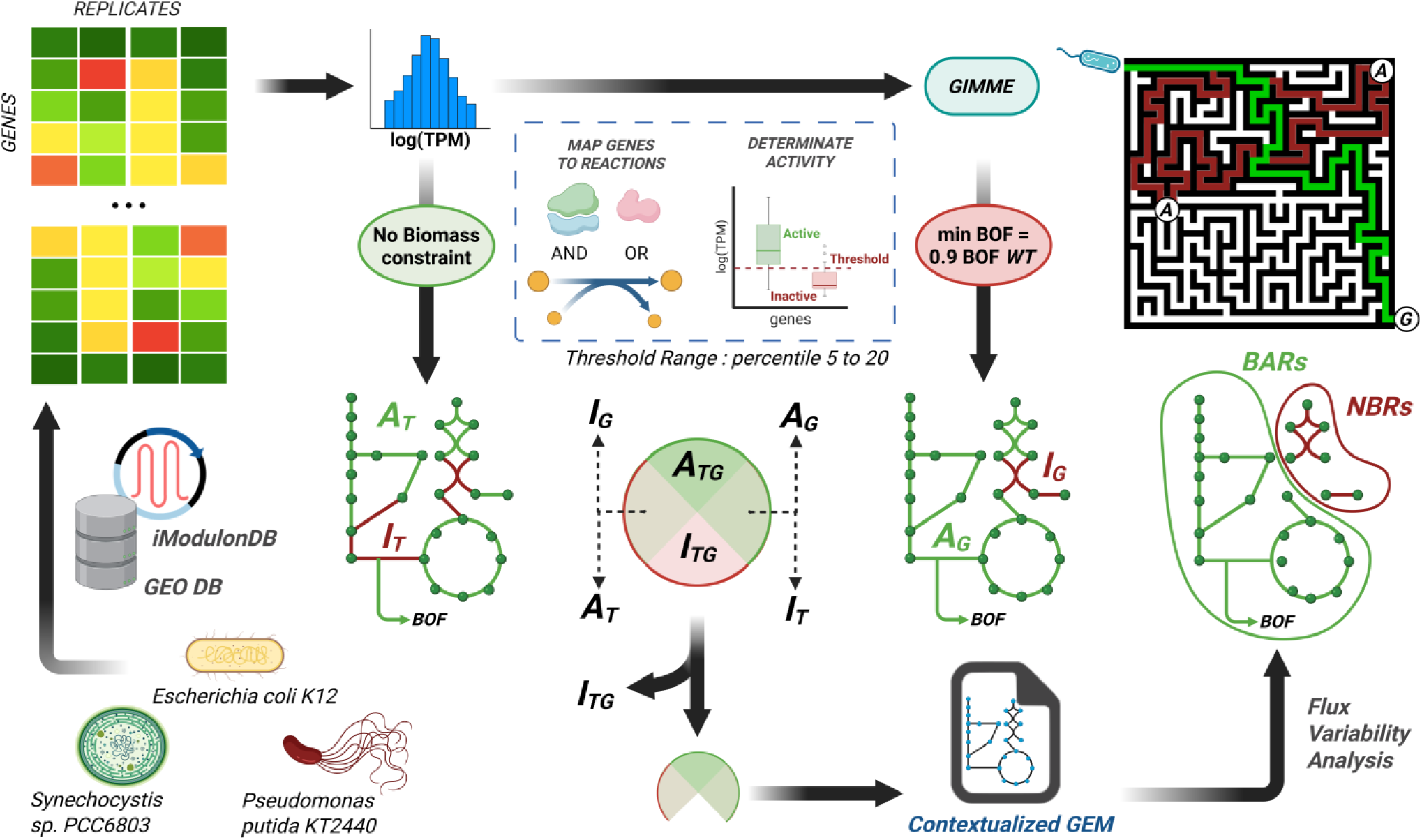
Schematic representation of EMBER workflow. The figure recapitulates the application of EMBER framework in this work. It starts with gathering transcriptomic data for diverse conditions from a source that equally pre-process sequencing data in order to minimize artifacts. Whit such data, models are contextualized considering or not a GEM objective for obtaining the active and inactive reaction sets for each situation (AG, AT, IG and IT respectively). The contextualized model is constructed with all reaction sets but the one formed by reactions inactive in both conditions (ITG). With it an FVA is applied to detect blocked reactions and thus differentiate between BARs and NBRs. On the upper-left side of the figure a maze is presented that synthesizes the conceptualization of BARs (green path) and NBRs (red paths). While BARs constitute a minimal reaction set for achieving growth (G), NBRs would provide additional paths providing additional functionality potentially enhancing the adaptability (A) of the organism.

Conceptually, reactions classified as BARs, would represent an optimal set of reactions for biomass generation in a given condition (illustrated as the green path in the maze, Fig. 11). In contrast, NBRs divert resources toward different objectives that, in turn, will provide adaptability to the system (depicted as the red paths in the maze). As result of that, the trade-off between growth and adaptability can be quantitively studied as the metabolic effort required for maintaining each reaction set. In accordance with this line of thinking, it can be hypothesized that organisms that prioritize adaptability over growth (i.e., organisms inhabiting unpredictable environments) should allocate a greater proportion of their resources to NBRs. Conversely, organisms that prioritize growth, which favor BAR sets, should channel more resources towards biomass generation.

As the approach described above is very sensitive to expression threshold values for determining reaction presence, EMBER do not perform a rigid classification based on a unique threshold value. Instead, the reaction classification approach is done for a range of thresholds varying from percentile 5 to 20. Then, reaction presence is computed as the ratio of the number of times it is present in a given set and the total number of thresholds tested. In this way, for a reaction to be included in one of the sets it needs to have a presence value above 50% of the tested percentiles. This pipeline is implemented in the Study-level_e-merlin_analysis.ipynb file of EMBER repository.

### System level analysis of adaptive response reaction subnetwork in bacteria

To gain further insight into the adaptability trade-off, a evaluation of this system-level feature was done in three biotechnologically relevant organisms from different ecological niches, specifically *E. coli*, *P. putida* and *Synechocystis* (see materials and methods for further details on organism selection). For each organism, we compiled a broad set of gene expression datasets encompassing environmental conditions representative of their respective ecological contexts (see Table 2). During the analysis, we realized that this step introduces potential biases into the analysis, as the selected conditions may differ in diversity or elicit stronger transcriptional responses across organisms. To mitigate this bias, we have considered the comparable number of conditions per organism (27 for E. coli and 34 for Synechocystis) and explicitly acknowledge its potential influence.

**Table 1.** Details of GEMs used in the present work.

| **Organism** | **Used GEM(s)** | **Reaction Number** | **Original publication** |
| --- | --- | --- | --- |
| *Escherichia coli* str. K-12 substr. MG1655 | *iJO1336* | 2583 | Ort et al. (2011)<sup>53</sup> |
| *Pseudomonas putida* KT2440 | *iJN1462* | 2927 | Nogales et al. (2020)<sup>57</sup> |
| *Synechocystis* sp. PCC 6803 | *iJN678* | 863 | Nogales et al. (2012)<sup>58</sup> |

**Table 2.**
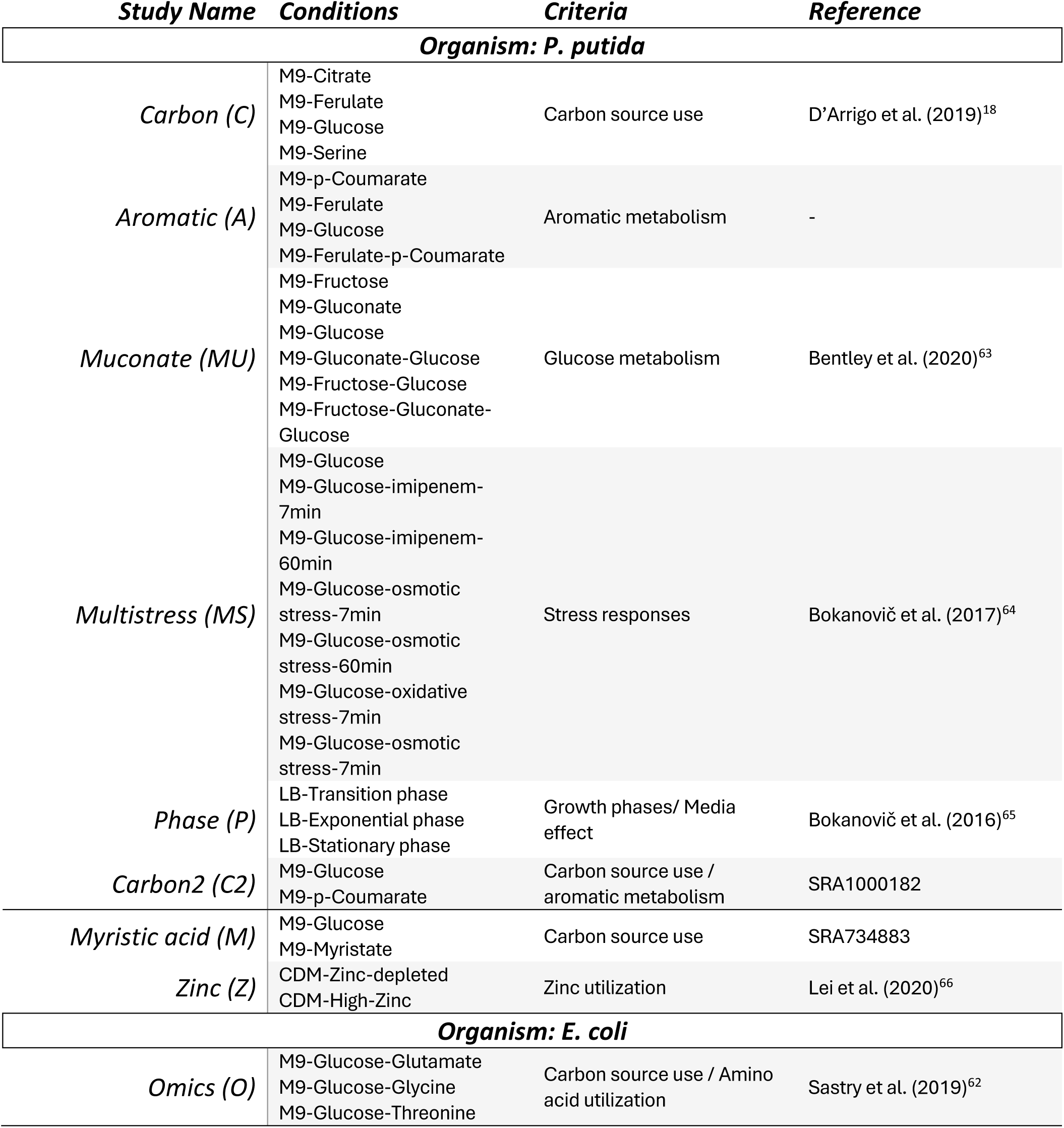
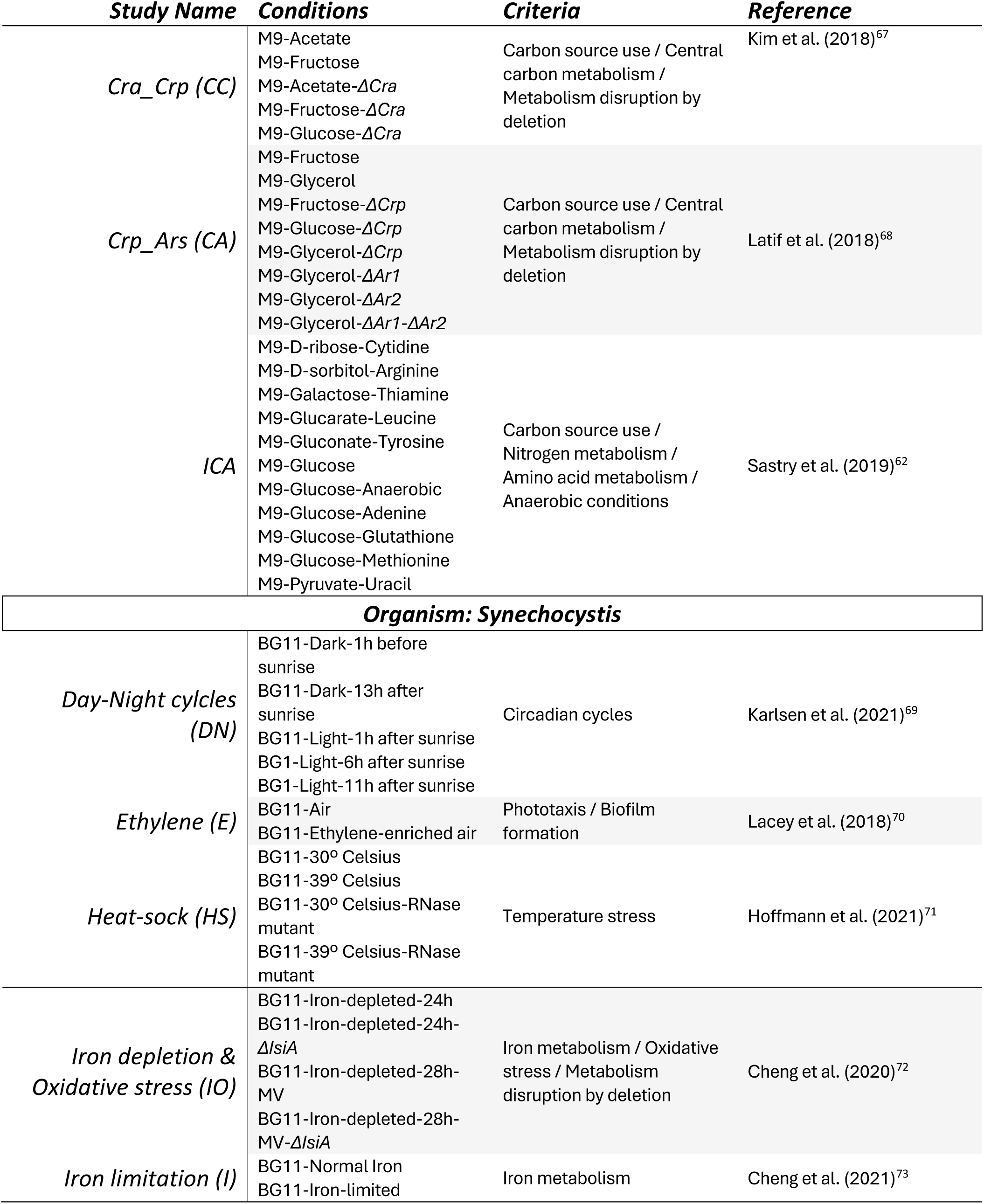
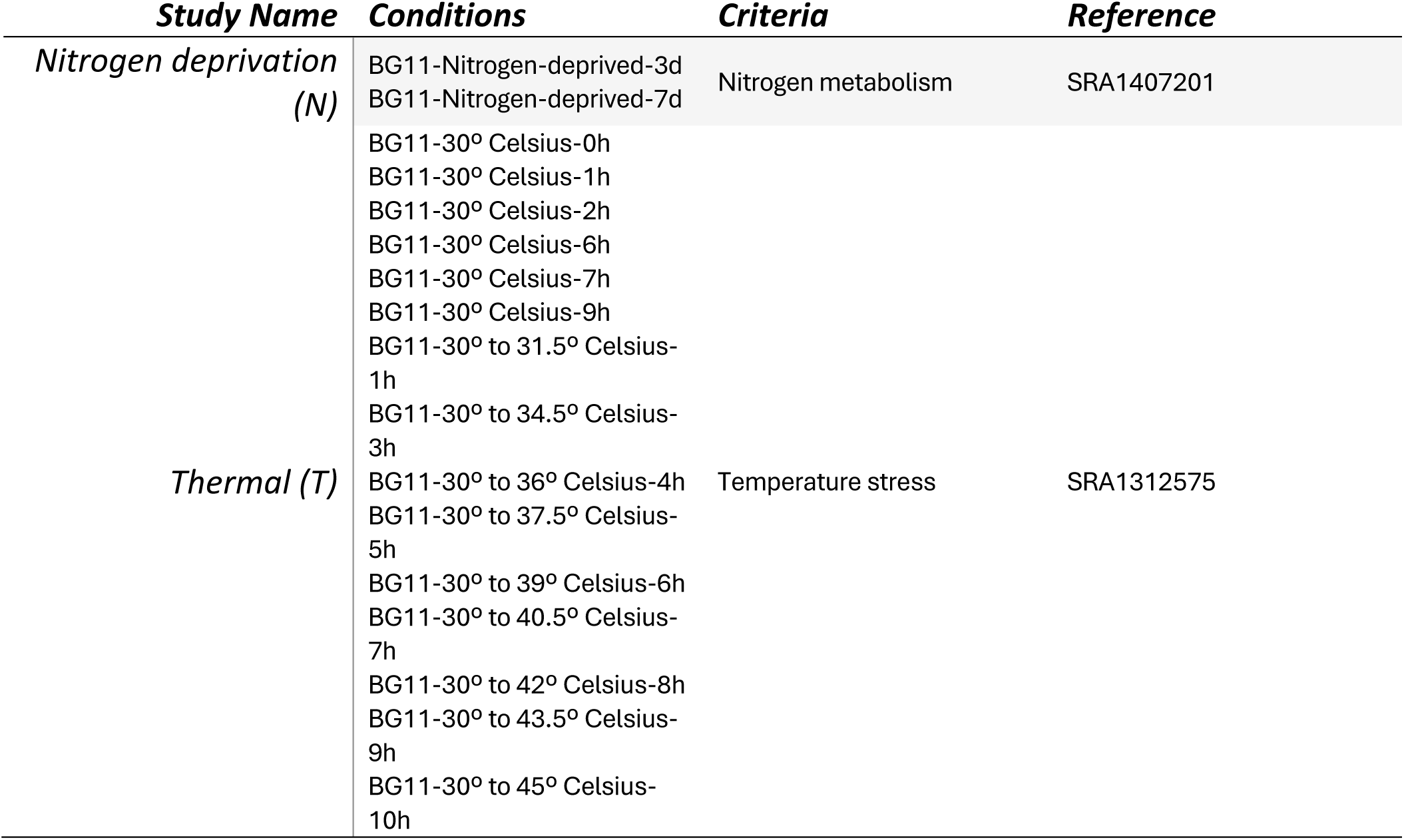
Conditions related to the selected transcriptomic experiments for each of the selected organisms. All datasets extracted from iModulonDB (for *P. putida* or *E. coli*) or generated with the iModulonDB pre-processing pipeline (for *Synechocystis*) are labelled under a given study name. Amon each of them, the inclusion interest criteria and refererences (when available) are shown.

The EMBER computational workflow classifies reactions as either NBRs or BARs. However, for translating this classification into a biologically interpretable framework, it was necessary to map reaction-level information to the corresponding genes, that will be further analyzed using several post analysis scripts (Figure 3). To perform the mapping itself, we applied the procedure detailed in *Gene classification* section of materials and methods. With it, we obtained the presence values of all genes classified as NBRs or BARs for each organism across the array of environmental conditions used in the analysis (Figure 4A).

**Figure 3.**
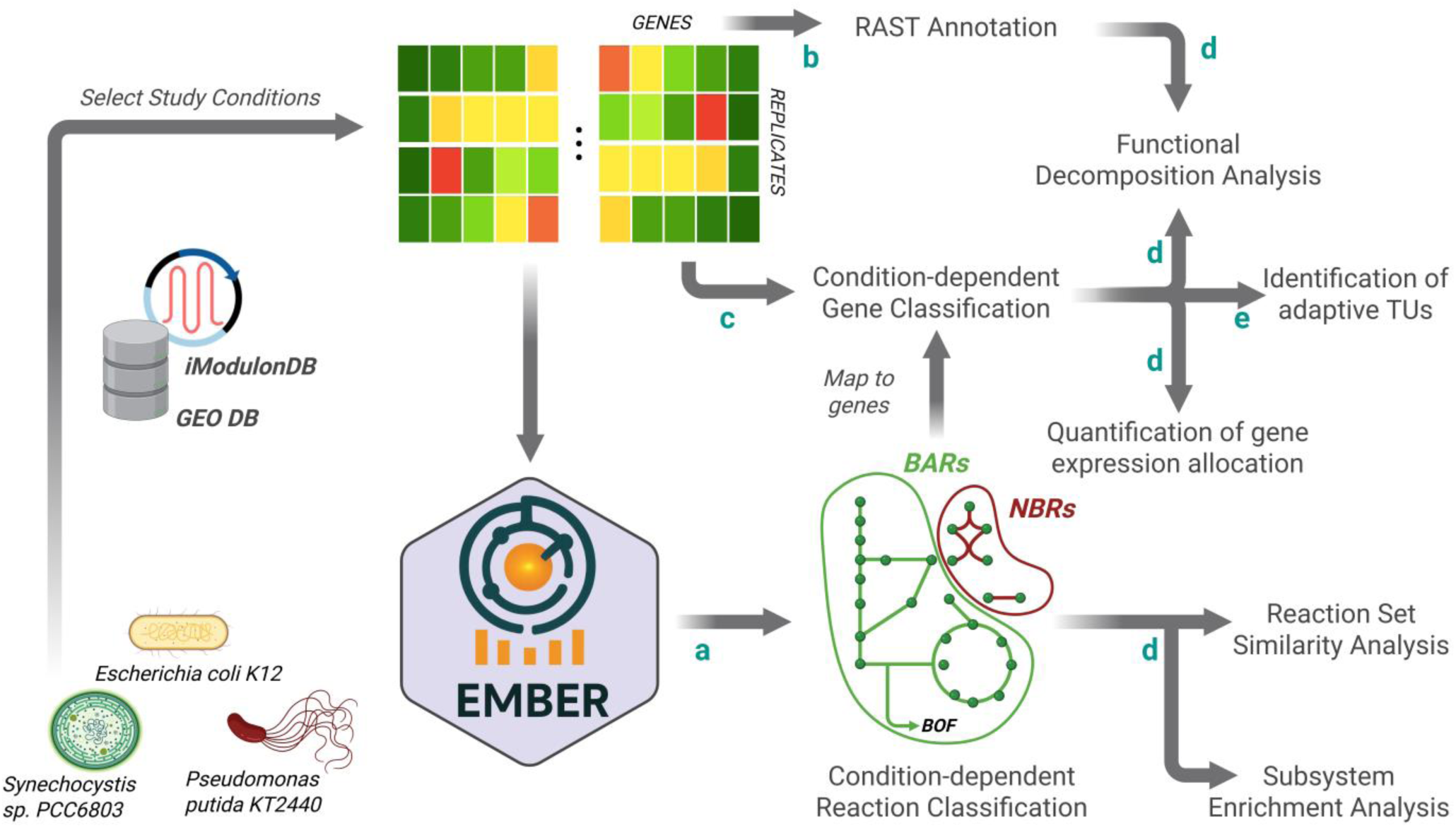
Integrated computational framework developed in this work to evaluate the growth-adaptability trade-off. The figure displays all the computational analysis steps involving the growth-adaptability trade-off made in this work. EMBER computes the condition-dependent classification of reactions in BARs and NBRs. Over the results obtained, direct analysis is made over the reaction sets for their quantitative characterization. Those sets are also mapped to the gene level for further analysis and the detection of metabolic engineering targets. Letters correspond to notebook files included in the EMBER repository implementing one or more analyses (**a:** *Study_level_e-merlin_analysis.ipynb;* **b:** *Functional_annotation_homogeneization.ipynb*; **c:** *Find_candidate_genes.ipynb;* **d:** *Comparative_Analysis.ipynb;* **e:** *Find_candidate_TUs.ipynb)*.

**Figure 4.**
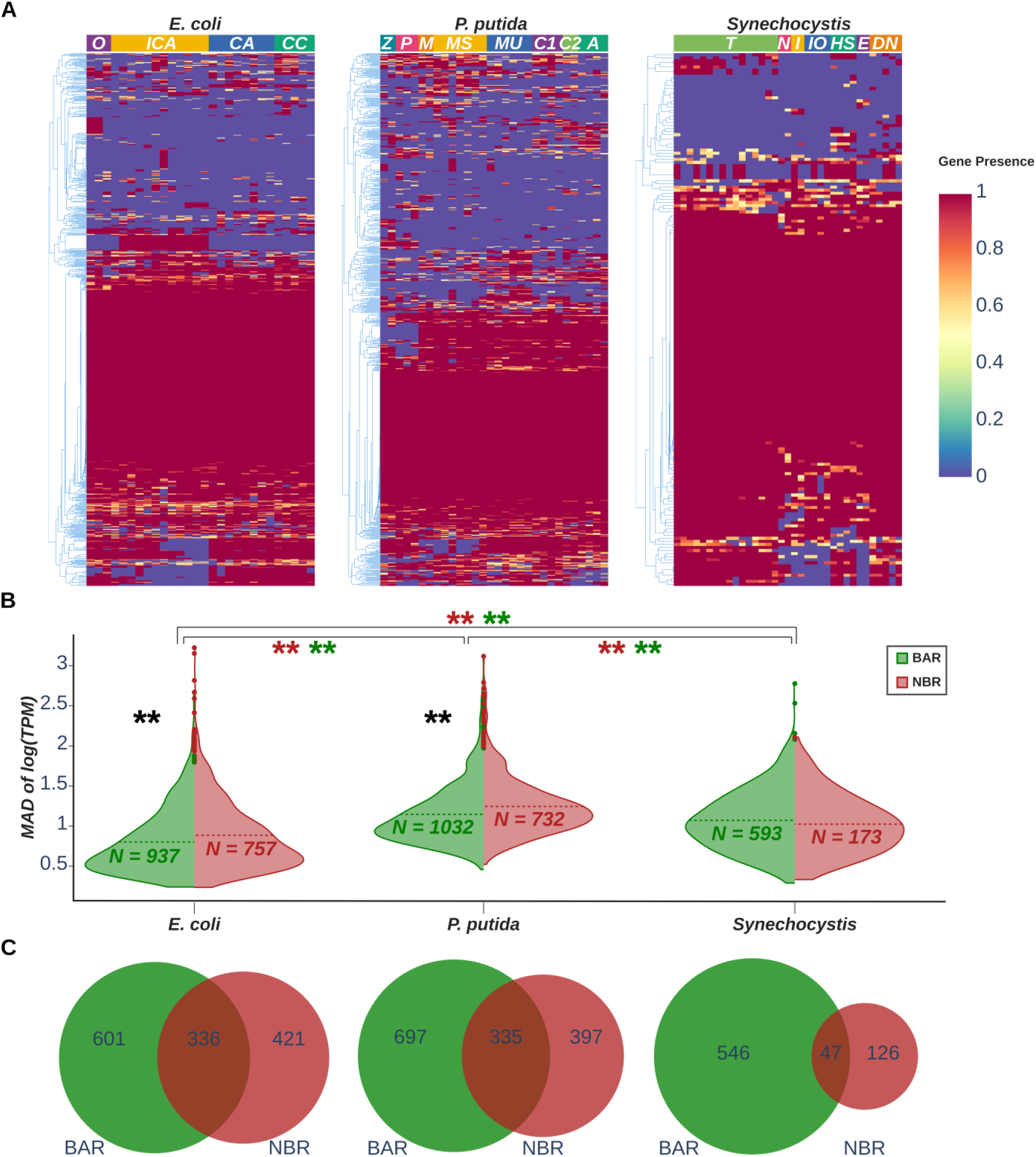
Classification and characterization of genes according to their role in adaptative responses. **(A)** Clustermaps showing the presence values of genes in NBR set computed according to section 0. Results are tagged with the abbreviations of their corresponding studies (see Table 2). **(B)** Violin plots summarizing the median absolute deviation (MAD) of all genes associated with NBR (firebrick) and BAR (green) sets. Double asterisks represent significance (p-value < 0.01) obtained by following a Mann-Whitney U test. The test was performed between data of same and different sets and organisms (as indicated by brackets), being the alternative hypothesis that the distribution with less median (dashed line) is stochastically less than the one being compared with. Also, for each distribution, the sample size is displayed in bold letters, being the numbers of genes that at least once belong to the correspondent set. (**C**) Venn diagrams proportionally displaying the genes always present in BAR (green) and NBR set (firebrick), alone with those present at least once in both (intersection).

An initial inspection of the NBR gene presence matrix across the three organisms revealed distinct topologies. Synechocystis exhibited a more homogeneous pattern, with a relatively consistent percentage of genes classified as NBRs across all conditions. In contrast, P. putida showed a more heterogeneous distribution, with a substantial fraction of genes classified as NBRs depending on the specific environmental condition. E. coli displayed an intermediate pattern (Figure 4A).

With respect to the absolute number of genes subject to study, *Synechocystis* emerged as the organism with the lowest number of transcriptionally active genes (766). This observation was attributed to the smaller gene content in the *Synechocystis* GEM, as well as to the relatively stable environmental conditions used in the analysis (e.g., autotrophic conditions). Notably, the proportion of genes consistently classified as NBR in *Synechocystis* was significantly lower compared to the other organisms (Figure 4C). Specifically, it exhibited only 17.5% of NBRs, in contrast to 27.8% and 31.0% in P. putida and E. coli, respectively. This suggests that *Synechocystis* allocates fewer resources to adaptative responses relative to the other 2 studied organisms, at least in the conditions analyzed.

We further focused on the gene expression variability across all replicate conditions tested in the transcriptomic datasets, analyzing both BARs and NBRs genes in the three species. This analysis revealed an interesting scenario (Figure 4B). The first notable observation was that, in *P. putida* and *E. coli*, genes likely associated with adaptability responses (NBRs) showed significantly higher expression variability compared to those involved in biomass and energy production (BARs), with p-values of 1.81·10-13 and 1.03·10-5, respectively.

This result further supports the hypothesis that NBR genes association with adaptability reflects the need for flexible expression patterns in such responses, whereas the biomass and energy production subnetwork is expected to maintain more stable expression levels. Interestingly, *Synechocystis* exhibited a similar distribution of expression variability across both gene sets, suggesting a more constrained regulatory architecture and, consequently, a rigid adaptability response compared to the other two organisms. Furthermore, beyond this data-driven observation, these results also contributed to deepening mechanistic understanding of adaptive responses.

As previously discussed, variability and stochasticity in gene expression can be used to distinguish between more passive adaptation strategies, such as BH from that more active responsive switching (RS) mechanisms^6^. Therefore, since significantly lower gene expression variability was observed in *E. coli*, it can be inferred that the adaptive response in this bacterium relies on RS rather than BH, in contrast to what may be suggested for *P. putida* (Figure 4B).

Taken together, these findings provide valuable insights into the adaptability strategies of the studied organisms. Both *E. coli* and *P. putida* appear to allocate substantial resources to adaptation. Based on gene expression variability, P. putida likely relies more on BH strategies, whereas E. coli seems to depend primarily on responsive switching RS mechanisms. On the other hand, *Synechocystis* appears to invest comparatively fewer resources in adaptability.

### Quantification of gene expression allocation reveals divergent adaptability strategies

In the previous section we estimated the global efforts towards biomass growth and adaptability of each organism based on the absolute number of genes classified as BAR and NBR, respectively. However, this metric does not capture the variability of such distribution across different conditions. To address this limitation, we developed a new approach, referred as adaptive effort analysis. This analysis involved mapping active genes corresponding to BAR or NBR for each condition tested in each organism. The NBR effort was then subsequently computed as the ratio of active NBR genes relative to the total active genes in each condition.

The results of this adaptive effort analysis was consistent with previous findings, preserving the relative ranking of organisms in terms of resources allocated to NBR functions (*Synechocystis* < *P. putida* < *E. coli*, Figure 5). However, distribution patterns reveal putative ecological contrasts: while *Synechocystis* exhibits an approximately normal distribution, *E. coli* and *P. putida* display profiles with broader ranges, particularly pronounced in *P. putida*. This suggests that adaptive responses in these heterotrophic bacteria are more variable and resource-intensive, reflecting ecological strategies that favor metabolic flexibility under fluctuating environments, whereas *Synechocystis* appears constrained by its photoautotrophic lifestyle.

**Figure 5.**
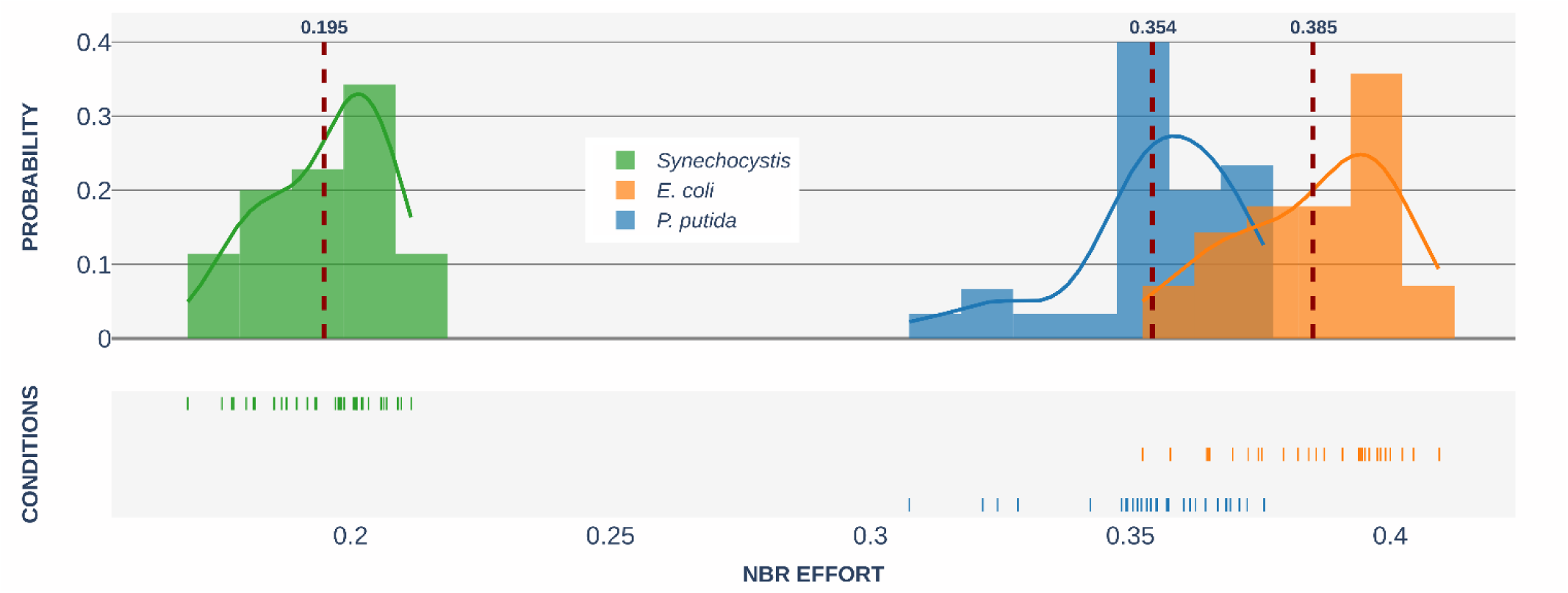
Results from the adaptive effort analysis (AEA). Histograms display the distribution of the NBR effort for each of selected organisms, computed as indicated in the present section. The means for each distribution are indicated by red dashed lines, with the actual value in bold letters above it. Below the histogram, the same data is represented through individual lines.

### Quantitative analysis of BAR and NBR sets similarity between conditions reveals organism-specific adaptive strategies

An important step in further characterizing the adaptability responses is to assess whether the computed sets of BAR and NBR reactions are similar between conditions or exhibit significant variation. Based on this rationale, it could be expected that BAR gene sets show lower variability between conditions, reflecting their role in core biomass production functions. Nevertheless, based on ribosome allocation, it would be expected to have a large impact of growth phase on variability, therefore, to ensure a fair comparison of innate metabolic responses, data from conditions involving genetic interventions (e.g. KOs) and growth conditions other than exponential growth phase were excluded from the analysis. The similarity of reaction sets within NBR and BAR categories was quantified using the Jaccard Index (JI). The code for this analysis can be found at the “Comparative_Analysis.ipynb” file within EMBER repository. The values of each pairwise comparison are shown in Figure 6.

**Figure 6.**
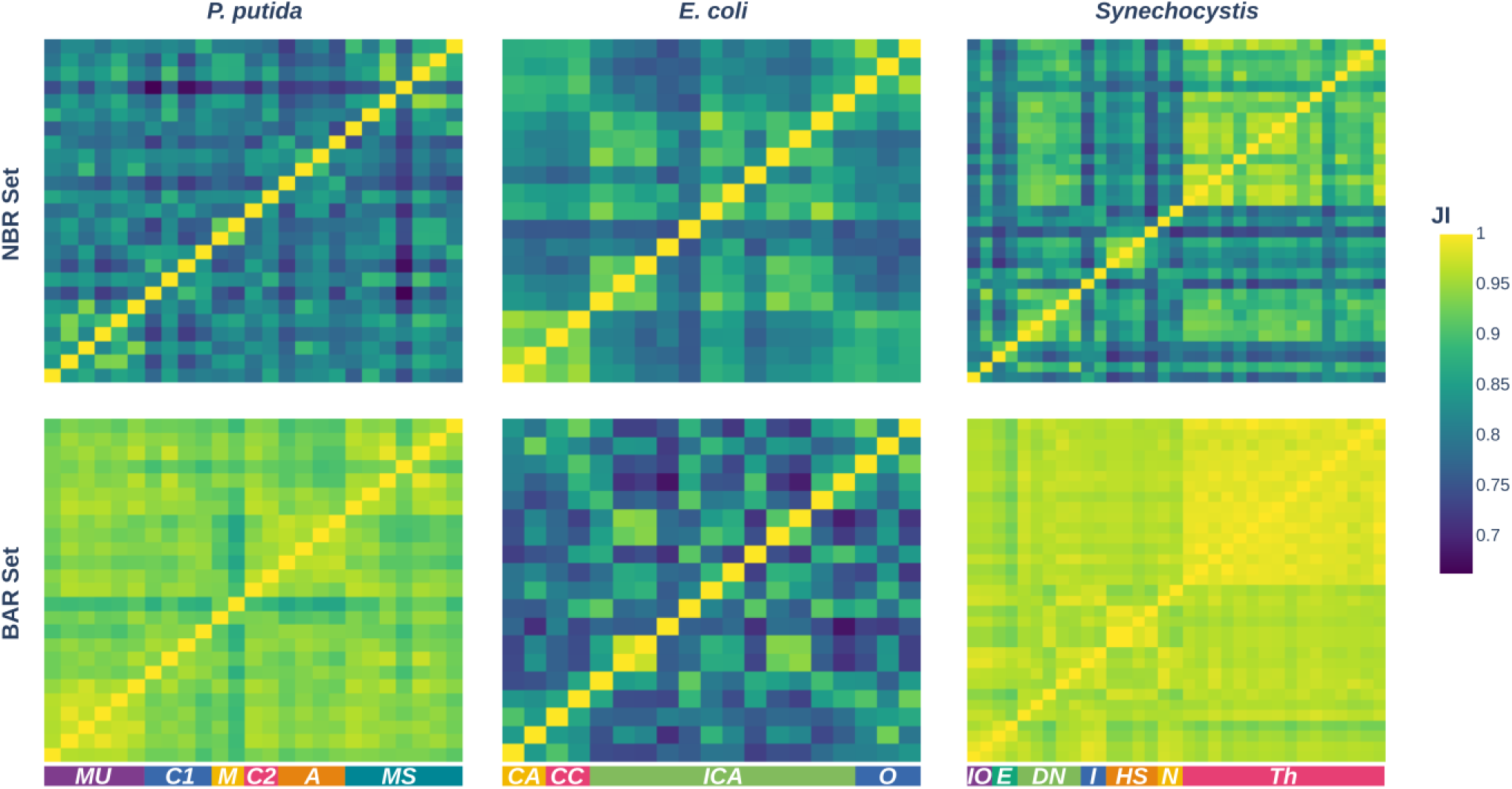
Results from the similarity analysis of reaction sets between conditions. Heatmaps showing JI values between sets of the same type and organism for all the studied conditions. Results are tagged with the abbreviations of their corresponding studies (see Table 2).

Interestingly, the analysis revealed that in *P. putida* and *Synechocystis*, the BAR set is highly homogeneous, with mean JI values between conditions of 0.94 and 0.97 respectively. *E. coli* exhibited a slightly more heterogeneous BAR set, with a mean JI of 0.80. Taken together, these results suggested that *E. coli* has a more diverse mode for biomass generation, in contrast to P. putida and *Synechocystis*, which maintain more consistent core biomass and energy production strategies. As expected, NBR behavior was more heterogeneous, apart from *E. coli*, which showed similar JI values for NBR and BAR sets (0.84 vs 0.80).

Notably, we observed modular patterns of JIs values for all sets of comparisons except for those involving NBR sets of *P. putida.* The patterns were primarily, though not exclusively, associated with specific study conditions, suggesting that each conditions elicit similar adaptive responses. Such consistency would be highly unlikely if the gene expression were driven mainly by stochastic noisy signals, thus fitting better with a regulated response such as the RS mechanism. From this perspective, all sets, except the NBR set in *P. putida,* appear to be influenced to varying degrees by RS responses. In contrast, the NBR set in *P. putida*, which also exhibits the lowest mean JI among NBR sets (0.81), is better explained by assuming that BH is the predominant driver of diversity between conditions.

Apart from all the observations above, this comparative analysis of adaptive and biomass-related responses across the three bacteria under study revealed organism-specific trade-offs shaped by distinct metabolic and regulatory architectures that will be thoroughly examined in the discussion section.

### Functional analysis of adaptive responses in bacteria

Once the quantitative characterization of adaptive responses was completed, we proceeded to assess its functional relevance in the three bacteria under study. This step is essential for biologically contextualizing the observed trade-offs and understanding the selective pressures that shape the regulatory and metabolic strategies of each organism in its native environment. Such functional insights are critical for guiding the condition-dependent selection of biotechnological targets, enabling more precise and context-aware engineering strategies.

To address the functional characterization of adaptive responses, two complementary classification systems were applied, each focused on a distinct biological entity—reactions and genes. The functional annotation of reactions was based on the author-defined subsystems provided within each GEM, allowing us to group reactions according to their metabolic roles. For gene-level classification, we extracted functional categories from the RAST server^17^, following the indications in materials and methods. This dual approach enabled a more comprehensive interpretation of the biological functions underlying the observed adaptive patterns.

### i. Enrichment analysis of subsystems of reactions

Assuming the aforementioned limitations, an enrichment analysis was performed for each of the studied organisms and conditions. This approach aimed to provide biological meaning to statistically derived BAR and NBR groups by testing whether they are disproportionately associated with known functional or regulatory categories. All the analyses were performed as described in materials and methods, obtaining the results showed in Figure 7. The corresponding results of the analysis are available in the “*condition-specific_reaction-enrichment_analysis.xlsx*” file within the results folder of each organism for the *EMBER* GitHub repository.

**Figure 7.**
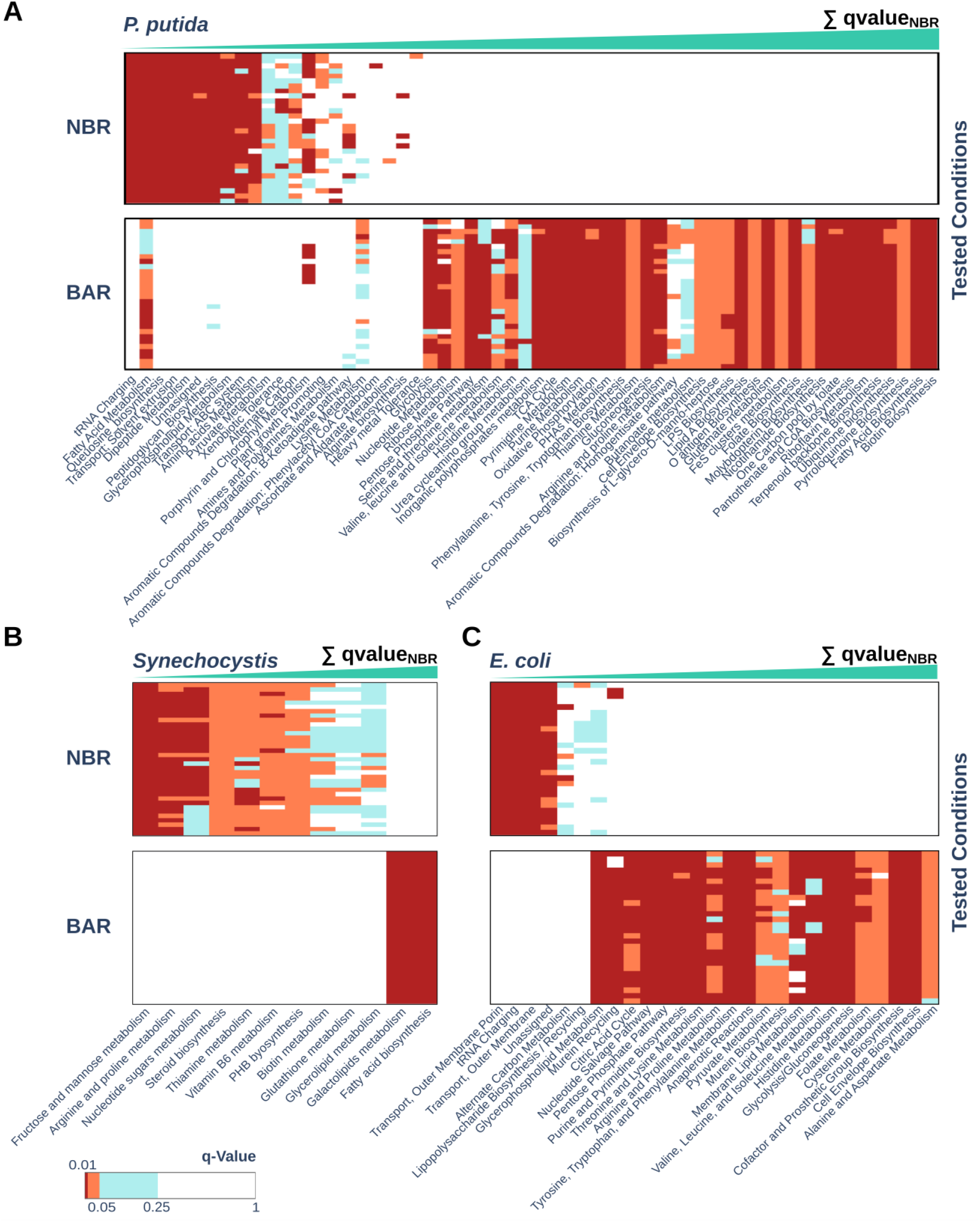
Enrichment analysis of reaction subsystems present in the GEMs of the selected organism. The figures show several heatmaps recapitulating results from enrichment analysis made for *P. putida* (A), *Synechocystis* (B) and *E. coli* (C) in each of their tested conditions and according to section 0. In the heatmaps, only subsystem found enriched at least once are shown. Those subsystems are sorted in descending order from left to right according to the total sum of their q-value in all conditions.

Despite direct comparison between organisms not being feasible due to annotation inconsistencies, all three models exhibit clear functional distinctions between their BAR and NBR reaction sets. Notably, the subsystems enriched within each category are largely orthogonal across conditions, enabling a robust functional characterization of adaptive responses.

Regarding *P. putida* BAR sets show a coherent functional composition, with enrichment in categories closely aligned with biomass production and core metabolic functions (Figure 7A). Notably, subsystems consistently enriched across conditions include canonical pathways of central metabolism such as the tricarboxylic acid (TCA) cycle, glycolysis, and the pentose phosphate pathway (PPP). Additionally, the biosynthesis of several amino acids and essential cofactors are also prominently represented. These findings support the notion that *P. putida* maintains a stable metabolic backbone dedicated to biomass generation in good agreement with the results from the similarity analysis (Figure 6).

Within the NBR sets of *P. putida*, transport-related subsystems—particularly those involving ABC transporters—emerged as some of the most consistently enriched categories (Figure 7A). This enrichment, together with the frequent presence of alternative carbon metabolism pathways across conditions in NBR sets, aligns with the well-documented adaptive strategy of *P. putida*, which relies on versatile transport systems to utilize a wide range of carbon sources^18^. Among the subsystems enriched in nearly all conditions, glycerophospholipid metabolism also stands out. This pathway plays a key role in fatty acid homeostasis and membrane remodeling, processes that are central to bacterial adaptation under environmental stress^19^. Dipeptide metabolism was also consistently enriched across all tested conditions within the NBR sets of *P. putida*. This aligns with the known ability of this strain to utilize dipeptides as a selective nitrogen source under nutrient-limited conditions^20^, a strategy that becomes particularly relevant in nitrogen-scarce environments, where specialized regulatory mechanisms have been proposed ^21^. Additionally, subsystems related to amine, amino acid, and polyamine metabolism—linked to alternative carbon and nitrogen sources—were enriched in fewer conditions, potentially contributing to this adaptive response.

Previously cited subsystems can be considered as default adaptive responses, as they are in almost all tested conditions. However, there are other interesting cases that appear to have random or condition-dependent patterns. An example of this are several subsystems that are composed of reactions involved in the aromatic compound’s degradation such as the xenobiotic tolerance (TNT degradation), the β-ketoadipate pathway and the phenylacetyl CoA catabolome. Interestingly, soil studies have shown that bacteria capable of degrading aromatic pollutants tend to persist and shape community interactions under contaminant pressure. Specifically, in soils amended with aromatic pollutants—such as polycyclic aromatic hydrocarbons, benzoates, chlorophenols, and related compounds—taxa including *Sphingomonas*, *Sphingobium*, *Achromobacter* and *Pseudomonas* are seen to emerge as dominant^22,23^.

Other relevant subsystems within this set consist of the plant growth promoting subsystem (harboring tryptamine metabolism reaction), which is enriched in stressful conditions like oxidative stress or zinc deprivation. Noteworthy, in this last condition, the heavy metal tolerance is also enriched. Finally, other interesting results consist of the enrichment of alginate biosynthesis in osmotic stress conditions (multi stress study and conditions with multiple carbon sources). As it is known, alginate takes part in biofilm formation responses, that could be a plausible adaptive response against stress conditions, including the osmotic stress^24^.

In summary, the functional landscape of *P. putida*’s NBR sets reveals a core of consistently enriched subsystems that underpin its adaptive versatility, complemented by condition-specific modules that reflect environmental sensing and stress resilience. This dual structure—combining default metabolic strategies with flexible, context-dependent responses—highlights the organism’s capacity to dynamically reconfigure its metabolic network in response to external challenges, reinforcing its ecological robustness and biotechnological potential.

Regarding *E. coli,* and as previously observed in *P. putida,* BAR sets also show coherent functional composition, with a high number of default-enriched subsystems linked to biomass-associated functions (Figure 7C). Among those, TCA, PPP, cell envelope biosynthesis, cofactor biosynthesis or nucleotide savage pathway clearly support that functional association. However, in terms of NBR results are much poorer than those obtained in *P. putida.* Although the two main transport subsystems are consistently enriched across conditions, suggesting similar adaptive responses, no other biologically meaningful subsystem follows this trend. Notably, one of the default enriched subsystems corresponds to unassigned functional reactions, limiting the biological significance of the analysis. These results, rather than reflecting a poor adaptive response of *E. coli*, likely stem from the incomplete and rather general definition of subsystems in the GEM of this organism. This translates into subsystems with a larger number of reactions, making enrichment both harder to detect and less informative—unlike in the *P. putida* GEM, where subsystem definitions are more precise and complete. We hypothesize that with a more comprehensive annotation, subsystems such as alternate carbon metabolism would show enrichment under a broader range of conditions.

Finally, results for *Synechocystis* were also quite limited (Figure 7B). BAR sets show coherent functional composition, dominated by biomass-associated functions like the fatty acid biosynthesis. Nevertheless, only 2 subsystems over the 55 present in *Synechocystis* GEM appear enriched in BAR sets. In the case of NBR, enriched subsystems included proline and arginine metabolism (*PLM*) and fructose and mannose metabolism (*FMM*), which could constitute important adaptive functions. However, the presence of cofactor biosynthesis complicates interpretation, and the overall number of enriched subsystems remains low. A few number of subsystems appear enriched in comparison with the total number of them. These limitations likely stem from the setup of the enrichment analysis, including subsystem size and the small BAR/NBR sets derived from *i*JN678.

Overall, it can be concluded that, despite obtaining promising results concerning the functional characterization of the adaptive responses in *P. putida,* results obtained for the other 2 organisms are more difficult to interpret. This led me to propose an alternative approach based on the previously classified genes, for which more standard annotation methods are available, leading to more informative and comparable results.

### ii. RAST functional categories composition analysis

The RAST server was employed further in this study to generate coherent and comparable annotations across the three selected species, enabling a functional analysis of the heterogeneity observed in metabolic adaptive responses. RAST offers several advantages over reaction-based subsystems. First, its annotations are designed for cross-species comparability, grounded in manually curated subsystems and FIGfams—protein families whose members are globally homologous and share identical functions—ensuring consistent functional assignments across organisms.

Moreover, RAST categories are hierarchically structured, allowing exploration from broad functional classes to more specific ones. This is particularly advantageous for gene set enrichment analysis (GSEA), where set size and specificity critically influence interpretability, as highlighted in the previous section. Finally, the RAST server benefits from continuous updates to the underlying SEED database, incorporating new subsystems and undergoing regular quality control, thereby enhancing annotation accuracy and consistency over time.

For taking advance of all these advantages, we generated the RAST annotations for the 3 selected organisms as indicated in section *Generation of homogeneous annotations with RAST* (Figure 8). All the generated code for this analysis can be found at the *“Functional_annotation_homogeneization.ipynb”* file within EMBER repository.

**Figure 8.**
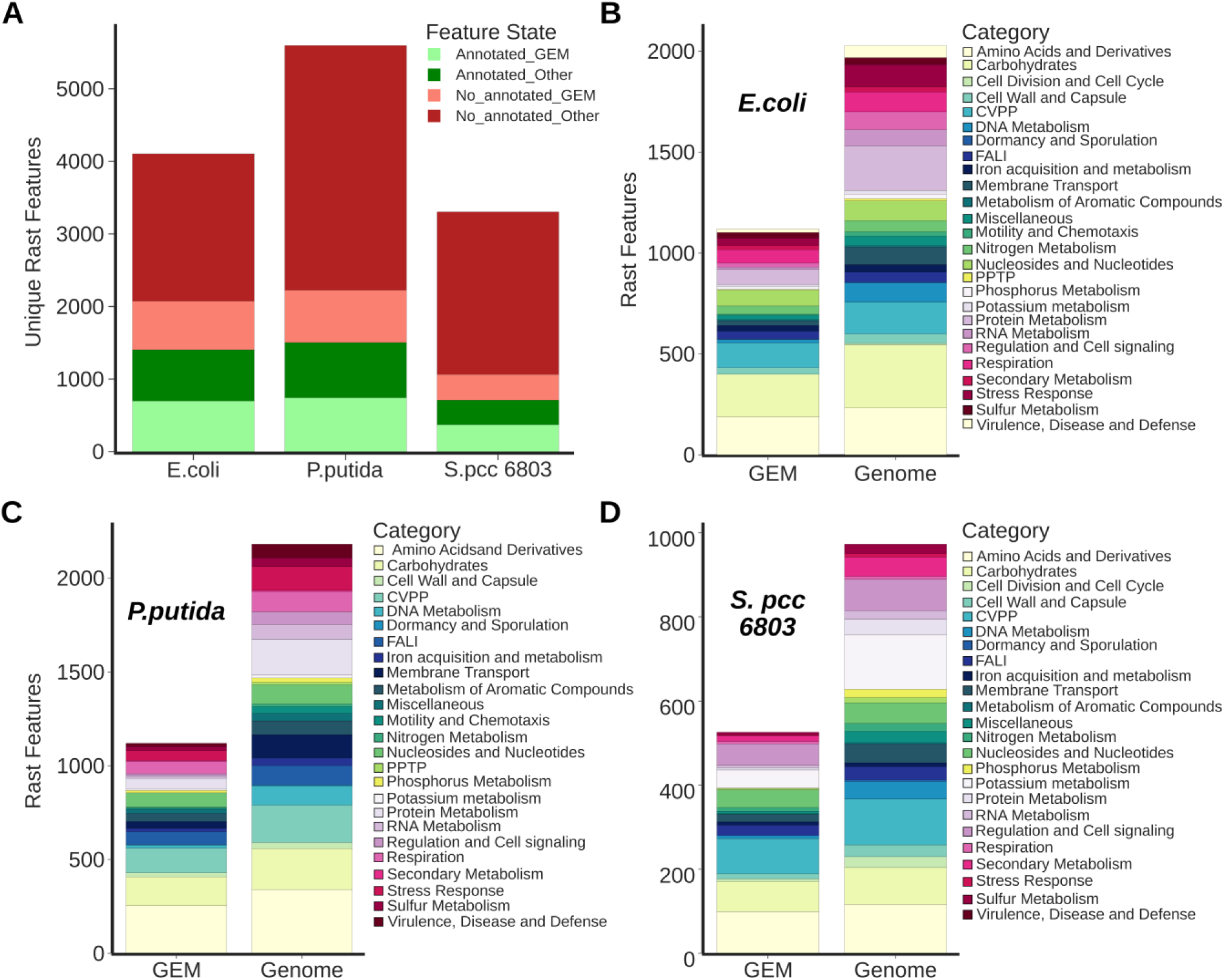
Summary of RAST annotation for the selected organisms. Histograms showing RAST annotation performance in terms of number of the unique features (FIGfams domains) that are annotated (**A**) and the categories obtained for *E. coli* (**B**), *P. putida* (**C**) and *Synechocystis* (**D**) are shown. Categories refer to the RAST functional class denominated as such, being the more general class generated by this program. Note that one same feature in RAST can be present in more than one category.

The annotations obtained comprise between 21.6% (*Synechocystis*) and 34.2% (*E. coli*) of total genome features with *P. putida* showing intermediate coverage at 26.9%. However, this percentage increases substantially when considering only GEM-associated features, with over half of them annotated—approximately 51% across all organisms. Those results are in line with the reported results on the original RAST paper^17^. Importantly, the annotations generated consistent RAST categories across all tested organisms (Figure 8B-D), with similar composition in both the GEM-associated and genome-wide features. This consistency thus enables a direct functional comparison of adaptive responses across species.

Functional analysis at the RAST category level revealed no major shifts in the composition of BAR genes across the studied organisms. In line with the subsystem-based analysis, *Synechocystis* displayed the most homogeneous behavior (Figure 9). Interestingly, some RAST categories appearing exclusively in BAR sets are shared across all three organisms. Notable examples include cofactor and prosthetic group biosynthesis (CVPP), fatty acid metabolism, and nucleotide/nucleoside metabolism—all closely linked to biomass maintenance. Their consistent presence suggests conserved core functions underpinning growth-related metabolic activity.

**Figure 9.**
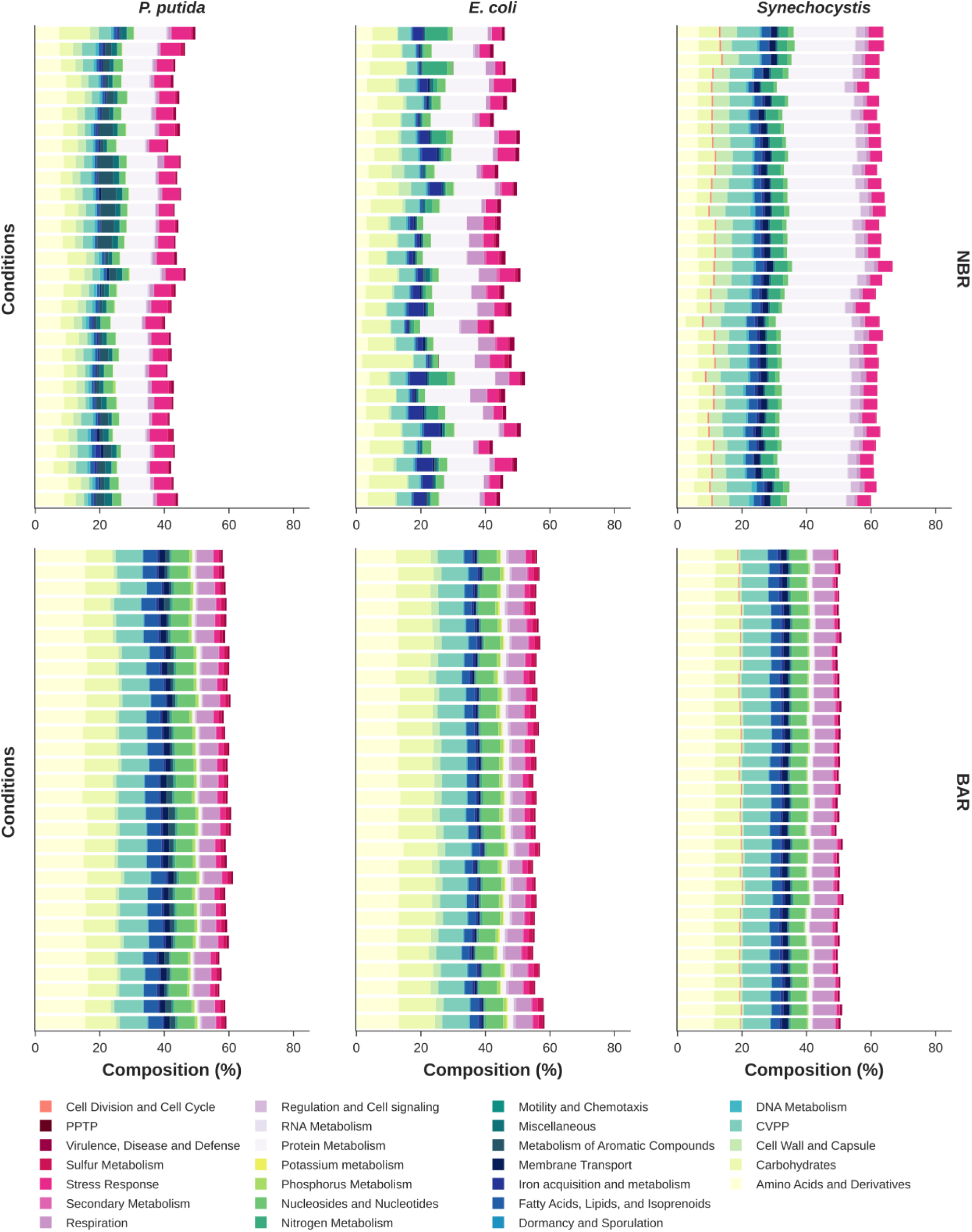
Reaction set composition across all tested conditions for each organism according to RAST categories. Histograms display the functional composition of BAR and NBR sets. Composition is represented in terms of percentage of all annotated genes for each organism, with colors representing different RAST categories. Abbreviations: PPTP (Phages, Prophages, Transposable elements, Plasmids), CVPP (Cofactors, Vitamins, Prosthetic Groups, Pigments).

In contrast, NBR sets showed considerable variation in functional composition across organisms (Figure 9). For instance, *P. putida* exhibited abundance in the metabolism of aromatic compounds, while *E. coli* showed it for nitrogen metabolism and iron acquisition. These categories were notably underrepresented in the respective BAR sets, including nitrogen metabolism in *Synechocystis*, where it appeared consistently across conditions. This pattern supports the adaptive role of these functions in response to environmental challenges. Additionally, some relevant functional patterns were conserved across species. Specifically, all NBR sets regardless of organism or condition showed enrichment in categories related to protein metabolism and stress responses, which were clearly underrepresented in the BAR sets. These functions may represent common adaptive mechanisms shared across organisms and warrant further investigation.

Although this preview offers some hints about the adaptive responses of the organisms, results are related to rather general functional systems. Thus, in order to deepen our understanding of adaptive functions we further subject the different annotation categories of RAST to an enrichment-based analysis.

### iii. Decomposition analysis of RAST functional categories

To comprehensively describe the functionality of the adaptive response we performed a decomposition analysis. This approach leverages the hierarchical structure of RAST annotations—comprising categories, subcategories, and subsystems, ordered from general to specific. Since each subsystem is nested within a subcategory, which in turn belongs to a broader category, this framework enables the assessment of functional responses across multiple levels of specificity, from high-level biological processes to fine-grained metabolic functions.

The decomposition analysis basically consists of computing the enrichment of each of those functional classes for each organism in each condition. Then, the enrichment frequency is computed relative to the number of conditions for each organism. Doing so, we can deconvolute the persistence of the metabolic responses for different specificity levels. This means that it is possible to track if general functions are coordinating more specific ones in a condition-dependent way or if their persistence is inherited from the persistence of a specific function. The code generated for performing this analysis is contained in the “*Comparative_Analysis.ipynb”* file within EMBER, with results represented in Figure 10.

**Figure 10.**
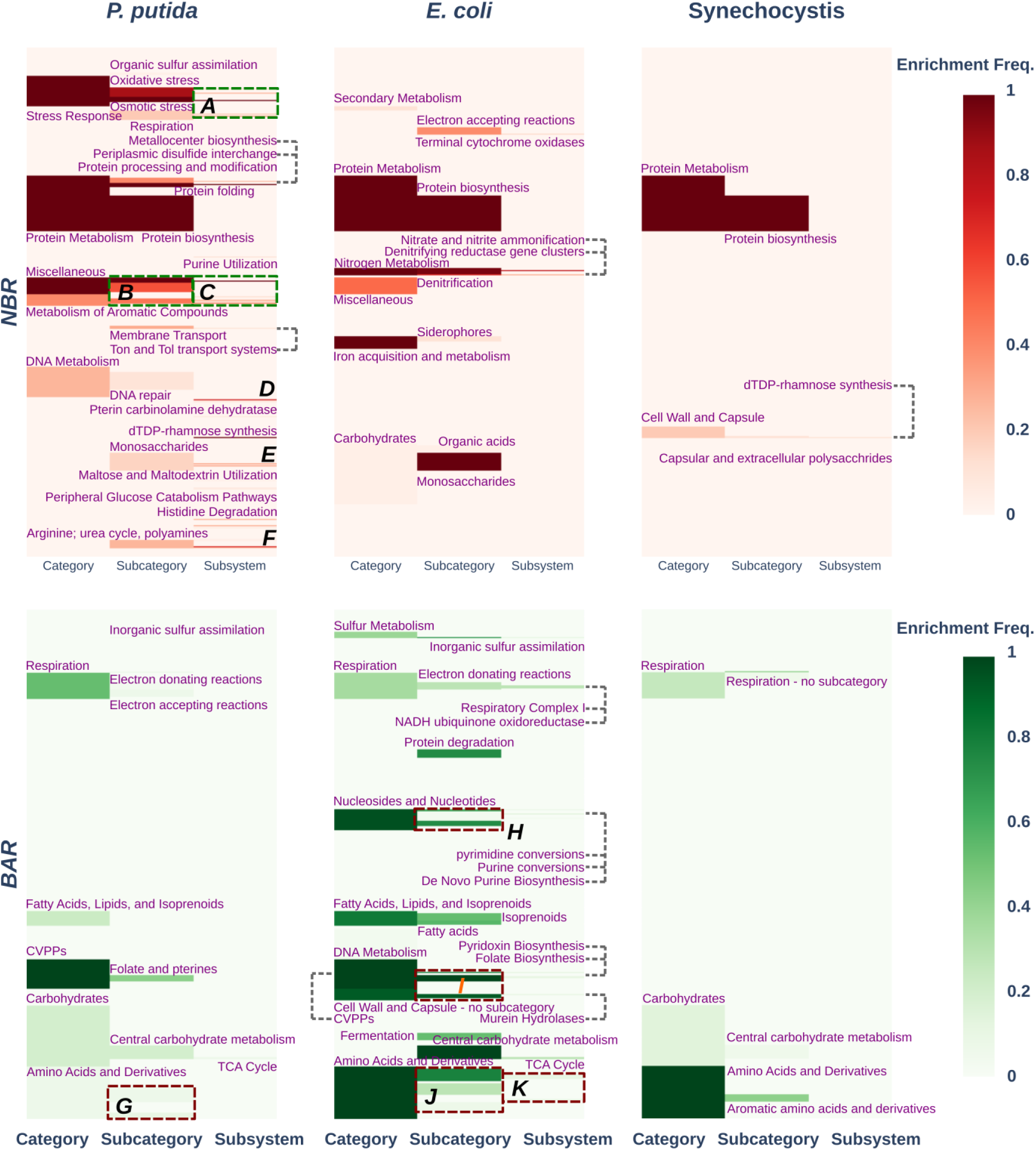
Results from functional decomposition analysis. Panel with heatmaps showing the enrichment frequency (percentage of conditions where q-value<0.05) of RAST functional classes among conditions used for each specie (columns) and reaction sets (rows; NBR in firebrick and BAR in green) used in this work. The size of each functional class is proportional to the number of annotated features that have been annotated with it. Note that distribution of those classes are always the same among the different heatmaps shown. Letters indicate groups of enriched classes that were too numerous to fit in the figure. (**A**) Glutathione: Non-redox reactions, Oxidative stress, Choline and Betaine Uptake and Betaine Biosynthesis, Formate hydrogenase and Biogenesis of c-type cytochromes. (**B**) Plant-Prokaryote DOE project, Miscellaneous and Metabolism of central aromatic intermediates. (**C**) Niacin-Choline transport and metabolism, Benzoate degradation, Catechol branch of β-ketoadipate pathway and Protocatechuate branch of β-ketoadipate pathway. (**D**) Cobalamin synthesis and Coenzyme B12 biosynthesis. (**E**) D-galactarate, D-glucarate and D-glycerate catabolism and D-gluconate and ketogluconates metabolism. (**F**) Leucine Degradation and HMG-CoA Metabolism, hydroxy-3-methylglutaryl CoA Synthesis, Aromatic amino acid degradation, Urea decomposition, Polyamine Metabolism and Urease subunits. (**G**) Branched-chain amino acids, Aromatic amino acids and derivatives and Alanine, serine, and glycine. (**H**) Pyrimidines, Purines and Nucleosides and Nucleotides. (I) Pyridoxine, Folate and pterines and Cell Wall and Capsule. (**J**) Lysine, threonine, methionine, and cysteine, Glutamine, glutamate, aspartate, asparagine; ammonia assimilation, Branched-chain amino acids and Arginine; urea cycle, polyamines. (**K**) Methionine Biosynthesis, Lysine Biosynthesis DAP Pathway, GJO scratch and Chorismate: Intermediate for synthesis of Tryptophan, PAPA antibiotics, PABA, 3-hydroxyanthranilate and more.

This analysis revealed several functional classes that are consistently enriched across species in both NBRs and BARs. For BARs, these functions primarily correspond to fundamental biomass-related processes, including respiration, central carbohydrate metabolism, amino acid and derivative metabolism, and fatty acid and lipid metabolism. In contrast, the NBR set shows enrichment in functions related to protein metabolism, with a particular emphasis on protein biosynthesis (Figure 10). Although protein synthesis is generally considered a biomass-related function, many NBR-classified reactions actually correspond to tRNA charging steps. These reactions are disconnected from the BOF in the GEM and therefore do not contribute to the model’s functional output. This explains why the protein biosynthesis category, dominated by amino acid–tRNA conjugation reactions, appears within the NBR set and why the tRNA charging subsystem is enriched. (Figure 7A, C).

*Synechocystis* exhibits limited functional annotation when using RAST, largely due to the high proportion of genes with unknown function and the comparatively smaller number of genes represented in its GEM. This constraint hampers statistical significance for most functional classes and reduces the interpretative value of the analysis. For example, the nitrogen metabolism category, which is ubiquitous in NBR set of *Synechocystis* (Figure 9), is not enriched in any condition. We detected enrichment in functional classes within the cell wall and capsule category. Specifically, this category was enriched under 20% of the tested conditions, with its most specific subclasses appearing in 11.4% of them. These functions involve dTDP-L-rhamnose biosynthesis, a metabolite essential for bacterial adaptability through the synthesis of surface polysaccharides. This compound plays a critical role in biofilm formation and architecture, influencing cell attachment and channel maintenance. Additionally, dTDP-L-rhamnose supports motility and mediates specific host interactions, which are key for survival across diverse ecological niches^25–27^.

Regardless, *E. coli* BAR decomposition analysis recapitulated at great extent enrichment results obtained with reaction subsystems. Likewise, it represents a clear focus with the central carbohydrate metabolism, cell envelope and nucleotide biosynthesis. Additionally, sulfur and amino acid metabolism appears enriched in BAR sets. In contrast, NBR analysis expanded previous findings, revealing three interrelated functional classes consistently enriched across most tested conditions: nitrogen metabolism, iron metabolism, and electron-accepting reactions related to respiration.

Nitrogen metabolism enrichment is mainly driven by denitrification and ammonification, with the denitrifying reductase cluster and nitrite/nitrate ammonification enriched in 28.6% and 78.6% of conditions, respectively. It’s well known that *E. coli* reduces nitrate to nitrite, providing an alternative respiratory pathway crucial for adaptation to oxygen-limited environments, with the latter also serving as a nitrogen source via ammonification^28^. This metabolic flexibility offers a significant fitness advantage in niches such as the urinary tract or inflamed gut, enabling proliferation and tissue colonization^6,29^.In line with this result, such response has been previously reported as a BH strategy in some other organisms^30^.

Another important adaptability function that is enriched by default in all conditions is the iron acquisition, with siderophore biosynthesis behaving as condition-specific response (enriched in 10.7% of conditions). This result is not a surprise, as Iron availability is a key driver of *E. coli* adaptive physiology, being required for respiration and DNA synthesis, therefore assuring its supply would be a reasonable adaptive strategy.

Additionally, another functional class enriched by default in all conditions is monosaccharides, that contain domains used for alternative carbon source utilization (including gluconate, fucose, galactarate, glucarate, galactonate, ascorbate, arabinose, galacturonate, glucuronate and xylose). Although the general rule in *E. coli* is that glucose represses alternative sugar systems via catabolite repression (low cAMP–CRP) and inducer exclusion, this repression is not absolute, with evidence showing that many systems “leak” or are partially active even when glucose is present^31–34^. Therefore, this result suggests that those systems follow a BH strategy that provides a clear adaptative advantage to *E. coli*.

For *P. putida*, decomposition analysis largely mirrors the functional enrichment obtained with reaction subsystems for BAR and NBR sets but provides a more detailed characterization of the latter. Specifically, for alternate nitrogen sources, several potential candidates emerged, including polyamine utilization (70%), purine utilization (10%), and urea utilization (23.3%). Regarding alternate carbon sources, the analysis highlights central aromatic intermediates and alternative sugars. Within the first group, benzoate degradation and the catechol and protocatechuate branches of the β-ketoadipate pathway stand out as condition-dependent candidates, enriched in 13.3%, 23.3%, and 46.7% of conditions, respectively. For alternative sugars, results point to the utilization of galactarate, glucarate, and glycerate (33.3%), along with peripheral glucose catabolism pathways (6.7%) and maltose/maltodextrin metabolism (3.3%).

An interesting case merging alternate carbon and nitrogen sources with stress responses is the niacin-choline transport and metabolism, which, like their parent classes (osmotic stress and miscellaneous respectively), is enriched in all conditions. It is well documented that Plants interact with bacteria through these compounds, so this result points to a codependence strategy between *P. putida* and the rhizosphere that will be further explained in the discussion section.

Another adaptive response that seems to be important in *P. putida* is the permeability of the membrane. This is observed through moderate enrichment frequencies in functional classes modulating it like the membrane transport (30%), which includes ABC domains and several metal cation-binding ones. Among the last ones are the previously mentioned Ton and Tol transport systems, essential for iron acquisition. Finally, another adaptability trait observed is the xenobiotic degradation response. This class is among 56.7% of conditions, including several dioxygenase domains, known for their role in the degradation of xenobiotic compounds.

Overall, when decomposition analysis detected enrichments, these largely recapitulated adaptive functions previously identified through the reaction subsystem approach, while providing higher resolution by linking overrepresented functions to the organism’s habitat and lifestyle. Notably, BAR and NBR sets were enriched in orthogonal functional classes, underscoring the growth–adaptability metabolic trade-off.

## 3. Discussion

### EMBER characterization of growth-adaptability trade-off recapitulates documented responses

Using EMBER, we uncovered a marked variability in metabolic efforts dedicated to maintaining adaptive functions across organisms. Specifically, *P. putida* and *E. coli* allocated more than one-third of their active genes to adaptive functions, whereas *Synechocystis* did not reach even one-fifth. Moreover, the reactions resulting from the expression of these genes were highly similar across conditions in *Synechocystis*, suggesting a more rigid metabolic architecture compared to the other two organisms. This observation points to fundamental differences in regulatory flexibility and metabolic plasticity among species.

Another relevant insight was that genes associated with adaptive functions (NBR set) exhibited greater expression variability compared to growth-associated genes (BAR set). This finding aligns with previous studies indicating that BH phenomena are largely mediated by stochasticity in gene expression^5,6^. For example, this relationship has been clearly demonstrated in *E. coli*, where noisy activation of the *gadBC* operon predicts single-cell survival under acid stress^9^. Notably, genes within this operon were among those detected by the approach developed in this work.

Interestingly, significant differences were observed in gene expression variability between the studied organisms for both BAR and NBR genes. While it is well established that species differ in their levels of expression variability^7^, the underlying causes remain an active area of basic research^1^. The analysis presented here advances this understanding by showing that variability is also linked to high-level functional roles of genes. Specifically, for NBR genes, higher expression variability suggests that adaptive responses rely more heavily on BH strategies, whereas lower variability indicates a stronger contribution from RS mechanisms. Therefore, the results obtained shed light on the general ecological strategies of the organisms studied.

For instance, *P. putida* exhibits significantly higher variability in both BAR and NBR genes, consistent with a strong reliance on BH strategies, while *E. coli* shows markedly lower variability in both sets, pointing to a predominant influence of RS (Figure 12B). From such discovery, new questions arise such as if such variability is a selected, organism-specific trait or, in contrary, is just a side effect of the complex interactions driving the genetic response of the organisms.

**Figure 11.**
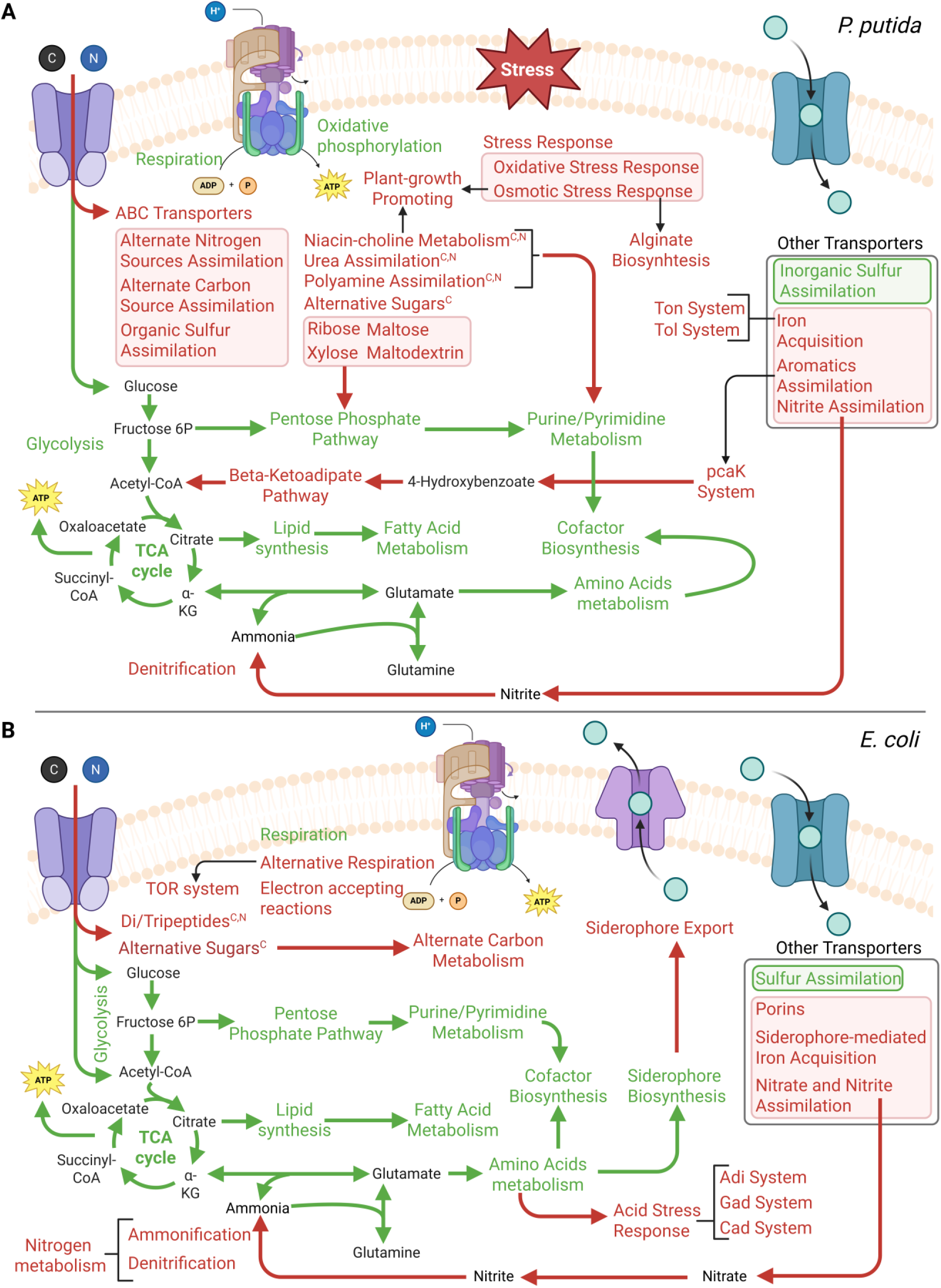
Schematic representation of results obtained through the application of EMBER enrichment post analysis. More prevalent enriched reaction subsystems and RAST categories for biomass (green) and adaptive functions are displayed in the metabolic context of *P. putida* (B) and *E. coli* (B). Superscripts refer to the identity of functions to provide either a source of carbon (C) or nitrogen (N).

**Figure 12.**
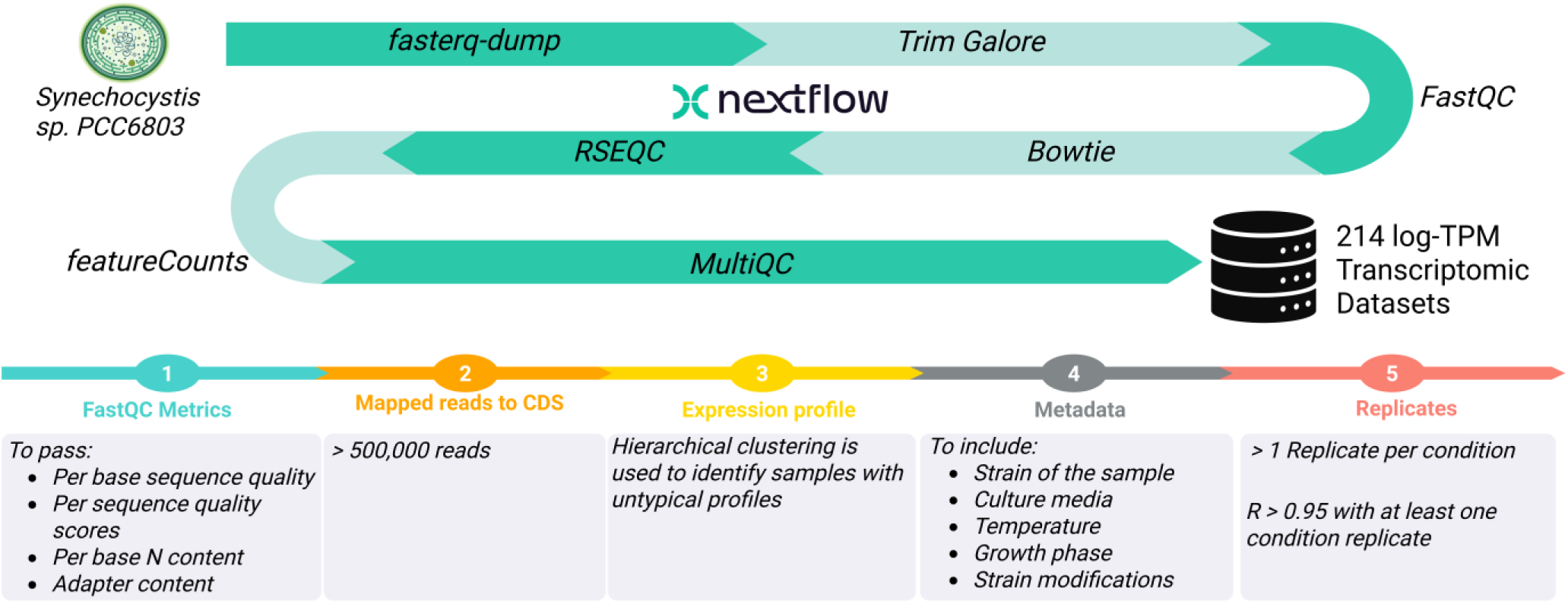
Schematic representation of the RNA-seq data extraction pipeline and quality filtering applied in this work.

The comparative analysis of adaptive and biomass-related responses across the three bacteria reveals organism-specific trade-offs shaped by distinct metabolic and regulatory architectures. Among them, *Synechocystis* exhibited the most homogeneous profile in both adaptability and biomass-associated responses, with gene expression patterns indicating a consistent and uniform implementation of RS. This resulted in low condition-dependent variability and a streamlined transcriptional landscape (Figure 6), supporting the idea that RS in *Synechocystis* operates in a highly coordinated manner to maintain functional stability across environmental conditions.

In contrast, *E. coli* displayed pronounced heterogeneity in its biomass-related metabolism, with variability levels comparable to those observed in its adaptive responses. This behavior is consistent with the presence of multiple metabolic modes for energy and biomass production, including aerobic respiration using oxygen as the terminal electron acceptor, anaerobic respiration relying on a diverse array of alternative electron acceptors, and fermentative metabolism in the absence of those external electron acceptors. The modular organization observed in both BAR and NBR responses strongly supports the role of RS in shaping the metabolic plasticity of *E. coli*, enabling transitions between distinct regulatory states in response to environmental cues.

Meanwhile, *P. putida* exhibited a relatively stable BAR profile, consistent with its obligate aerobic metabolism, while its adaptive responses were markedly heterogeneous. Notably, the absence of modular patterns linked to environmental conditions and the lack of evidence for RS suggest that adaptability in *P. putida* is primarily governed by BH strategies. These are characterized by distributed, context-specific regulatory mechanisms that enable flexible, non-modular responses to environmental perturbations. Collectively, these findings underscore the organism-specific nature of adaptive trade-offs and highlight the interplay between metabolic versatility, response dynamics, and regulatory architecture.

Finally, functional characterization efforts of all extracted sets reveal that both sets recapitulate coherently traits associated with the goals of adaptation or growth. In that sense, while BAR sets are enriched in biosynthetic functions, NBR sets reflect responses to several types of perturbations or stresses. Moreover, those adaptation responses closely link with the native environment of the organism.

For example, *P. putida* showed enrichment in transport systems that enhance membrane permeability, pathways for aromatic compound degradation, and osmotic stress responses—features consistent with metabolic challenges commonly encountered in soil environments (Figure 11). Additionally, the enrichment of niacin and choline metabolism within the NBR set illustrates adaptive strategies involving inter-species cooperation in the rhizosphere. It is well documented that plants under water or osmotic stress exude choline and glycine betaine, which bacteria—including pseudomonads—import and convert into osmo-protectants that improve survival and root colonization, or into metabolic intermediates readily used as carbon and nitrogen source^35,36^. Similarly, microbial uptake and metabolism of niacin (a precursor of NAD) can influence NAD availability in the rhizosphere, indirectly affecting plant redox balance and immune signaling, and promoting a shift from growth to defense metabolism^37^. Together, choline and niacin transport shape microbial fitness, community composition, and ultimately plant resilience, acting as metabolic links in cross-kingdom communication. The fact that both systems consistently appear within the NBR set under all conditions suggests a clear BH strategy to mitigate the risks posed by unpredictable changes in the rhizosphere—a habitat well known to *P. putida*.

*E. coli,* on the other hand, prioritizes respiration adaptability or the acquisition of nitrogen or metal ions (Figure 11), with the case of iron acquisition being very informative. This metal is scarce in many of *E.coli* natural niches due to its low solubility and host sequestration in mammal gastrointestinal tract^38^. To cope with it, *E. coli* deploys a redundant toolkit—high-affinity siderophores (like enterobactin, salmochelin or aerobactin), heme and ferric-citrate uptake, and ferrous transport systems—plus TonB-dependent importers to capture iron from diverse sources^39,40^. Also, it has been shown that regulation by Fur integrates iron uptake with oxidative-stress and metabolic responses, minimizing toxic free iron while enabling colonization and competitive fitness in host and environmental settings^29,41^. Therefore, the enrichment of this subsystem in the NBR set makes perfect sense considering *E. coli* biological context, where oxygen fluctuates and nutrient levels are compromised due to competition with other bacteria, such as the gastrointestinal tract of mammals^38^.

Despite these findings, functional characterization results for *Synechocystis* were very limited. This limitation is likely due to several factors. First, the GEM representing this organism contains considerably fewer genes than the other two models, which reduces the size of both BAR and NBR sets. A similar constraint applies to the number of RAST annotations available. Altogether, these factors hinder the detection of statistically enriched functional categories, making the results for this organism less informative.

Such limitations are expected for poorly characterized strains with scarce annotation data or for organisms whose sequences exhibit low homology values, complicating functional assignments. In this context, novel annotation approaches that do not rely on homology could significantly improve the functional characterization of non-model organisms. Specifically, methods based on protein language models have shown promising results, achieving higher annotation coverage even for specific GO terms^42^. Therefore, a potential improvement to the present analysis would involve applying these models to densely annotate the selected organisms, thereby enhancing the depth and reliability of functional insights.

### Integrating the reshaping of different metabolic trade-offs for bioprocess optimization

Assuming the multioptimality hypothesis, bacterial metabolism balances several biological trade-offs, promoting metabolic heterogeneity via BH or RS. Those trade-offs arise from stoichiometric, enzymatic, and resource constraints, requiring compromises among growth, energy efficiency, stress resistance, and adaptability, with the present work focusing on the latest.

An important question arising from this trade-off is whether the adaptive response constitutes a universal and homogeneous phenomenon across the population or, alternatively, operates at the single-cell level, exhibiting pronounced cell-to-cell heterogeneity. In this regard, the developed approach has a fundamental limitation. This consists of the impossibility of differentiating if a response is taking place in the whole bacterial population or just in a small fraction of it. Moreover, it is likely that subpopulation responses are underrepresented in this approach due to lower levels of transcripts of these responses in bulk RNA-seq experiments in comparison with whole-population responses.

Beyond this conceptual inquiry, a derivate of this work is to exploit these mechanisms to enhance biotechnological performance of bacterial strains. As previously mentioned, potential strategies include emulating division of labor or eliminating default adaptive genes that confer no advantage under biotechnologically relevant conditions, thereby reallocating resources toward the desired application.

Moreover, the engineering of this trade-off could be applied additively with a widely used one, the growth-coupled (GC) phenotype. This metabolic state consists of a readjusting of the growth-production trade-off, referring to a metabolic configuration in which cell growth is inherently linked to the synthesis of a target metabolite. In such systems, the production of the desired compound becomes a prerequisite for cellular proliferation, effectively aligning evolutionary pressures with biotechnological goals^43,44^.

The simultaneous application of both approaches, at least theoretically, should not concern any problem aside from the potential metabolic burden we are introducing the organisms involved. For example, by deleting some default, redundant systems that generate unnecessary metabolic cost in a bioreactor set up, we could compensate for the reduction in growth often generated by GC phenotypes.

Another possibility, although more elaborate one, could consist of using a promoter candidate that mediates DoL for distributing the pathway of a complex product for which a GC design has previously been found. In this latter application, once the design of a GC phenotype, including the heterologous reactions, is reached computationally, the enzymes participating in the pathway should be subjected under the expression control of a DoL promoter that could be either a default one or a conditional, depending on the specific bioprocess.

Importantly, many computational methods exist for the redesign of the growth-production trade-off through GC phenotype^44^, contrary to the growth-adaptability one. Therefore, to advance on the described integration, differentiating between types of adaptive responses will be crucial for the final application of a given candidate, with the experimental validation of our model-driven being essential. This could be done, for example, by mapping the EMBER results to transcriptional units (TUs). Specifically, their promoters could be included in the strain design phase, implementing one of the biotechnological tasks by harnessing this trade-off. In that way, available databases such RegulonDB^45^ could be integrated with EMBER for providing a new tool in metabolic engineering that redirects the adaptive programming of bacteria towards complex, biotechnologically relevant tasks.

## 4. Materials and Methods

### Selection of chassis and their computational representation

The selection of specific bacterial strains for investigating metabolic heterogeneity through genome-scale metabolic models (GEMs) is crucial for leveraging existing knowledge and experimental capabilities. In the present study, we have chosen 3 distinct organisms: *Pseudomonas putida* KT2440 (*P. putida*), *Escherichia coli* str. K-12 substr. MG1655 (*E. coli*) and *Synechocystis sp.* PCC 6803 (*Synechocystis*).

I thought these organisms represent ideal models for studying metabolic heterogeneity due to its presence in different ecological niches with substantial differences in composition and dynamical changes. Importantly, they possess well-validated GEMs (see Table 1), that is a necessary requirement for the generation of trustful results. In the following paragraph, some characteristics justifying this selection will be discussed:

- ***P. putida*.** This organism is a soil-dwelling bacterium. The unpredictable changes that occur in this habitat greatly contributes to is known robust metabolism. It has a primarily heterotrophic lifestyle, showing remarkable metabolic versatility and stress tolerance, including resistance to heavy metals and organic solvents. In this way, it is capable of aromatic compound degradation and lignin metabolism^46–48^. Importantly, it has been shown that it can survive under various environmental conditions including limited nitrogen^49^. Those characteristics have encouraged its use as synthetic biology chassis, enabling the characterization of some mechanisms underlying its intracellular heterogeneity^50,51^. Among those efforts, the generation of several high-quality GEMs of this organism is crucial for my approach, as it provides a mechanistic template where the characterization of its metabolic heterogeneity can be thoroughly analyzed. All the mentioned characteristics made *P. putida* the model organism used along all sections of this thesis.
- ***E. coli.*** This organism is commonly found in the digestive system of humans or animals. Consequently, *E. coli* is constantly exposed to extreme acid stress. Not surprisingly, studies have revealed significant heterogeneity and division of labor in *E. coli*’s acid stress responses, such as the one mediated by its Gad system, crucial for *E.coli* survival^9^. Also, such habitat also favors the adaptability of this organism to both aerobic and anaerobic conditions and a diverse carbon source utilization^6^. This heterogeneity extends to various metabolic activities, where individual cells exhibit diverse behaviors, even under well-controlled conditions^9,52^. For example, studies have observed heterogeneity in respiratory activity and lag time among *E. coli* colonies, suggesting that different metabolic strategies can be equally optimal^5^. Furthermore, research has identified metabolic coordination within *E. coli* biofilms, involving cross-regional resource allocation, local cycling, and feedback signaling, supported by spatially specific activation of metabolic pathways and strengthened transmembrane transport^8^. Importantly, *E. coli* is widely regarded as a model organism in microbiology and metabolic engineering, benefiting from an extensive body of accumulated knowledge and genetic tools. Proof of this is the fact that *E. coli* has several available GEMs, having 6 for the specific substrain selected. This further provide a robust foundation for computational analyses of metabolic fluxes and network operations^53,54^.
- ***Synechocystis***. This freshwater cyanobacterium is the most comprehensively studied organism of its phylum. It is a key organism in global ecosystems, being one of the few prokaryotes capable of oxygenic photosynthesis. This enables it to play a significant role in global carbon fixation and marine nitrogen fixation. Thus, its selection is crucial for extending heterogeneity studies to phototrophic organisms, offering a distinct metabolic framework compared to heterotrophs like *E. coli* and *P. putida*. Moreover, this organism exhibits metabolic flexibility as it has metabolic modes enabling growth on heterotrophic, photoautotrophic and mixotrophic conditions. It is adapted to diurnal cycles with distinct light and dark phase metabolism. While approximately 30% of its coding sequences have assigned functions, predominantly based on homology with genes from other bacteria, ongoing research aims to experimentally validate these functions and tools are in development to accelerate characterization efforts. The organism possesses a unique ultrastructure, including internal thylakoid membranes, which necessitate specialized cellular systems for targeting proteins and metabolites. This complex internal compartmentalization suggests additional layers of spatial and metabolic heterogeneity that could be investigated. Finally, from a synthetic biology perspective, *Synechocystis* is a promising chassis as it can produce interesting plant-derived compounds such as terpenes. Furthermore, it holds the potential for bacterial biofuel generation^55^. However, nowadays this bioprocess is still considered unfeasible as the energy return on energy invested (EROEI) is unfavorable, being an active research field^56^.

### Data acquisition

As previously mentioned, to enhance the predictive capabilities of genome-scale metabolic models (GEMs) and to accurately capture dynamic cellular behaviors, the integration of high-throughput omics data, particularly RNA sequencing (RNA-seq), is essential^59,60^. This addresses the inherent limitations of purely stoichiometric GEMs, which often lack the detailed context of gene expression changes influencing metabolic flux^16^.

In this light, the iModulonDB (https://imodulondb.org/) has proven to be a crucial resource, as it provides a quantitative framework for understanding microbial transcriptomic data, enabling its comprehensive interpretation. The content of iModulonDB is generated through a rigorous pre-processing pipeline for RNA-seq data, which is explained in the following section. The data resulting from this pipeline is processed through Independent Component Analysis (ICA), an unsupervised machine learning algorithm, is applied to the high-quality expression compendium to identify statistically independent components, which are subsequently transformed into iModulons via a thresholding process. Each iModulon within the database is a quantitative representation of a regulatory module, comprising gene weights for every gene and activity level for each experimental condition. These activity levels reflect the overall state of a transcriptional regulator across environmental conditions. This detailed characterization allows for biological interpretation and the discovery of novel regulons. This approach, demonstrated to outperform other regulatory module detection algorithms, ensures the identified modules are robust and biologically meaningful representations of the transcriptome^61^.

Access to iModulonDB is provided via its interactive website, which includes searchable dashboards for each iModulon. Additionally, the PyModulon Python package provides a programmatic interface for *i*Modulon analysis, characterization, interpretation, and visualization, allowing researchers to explore *i*Modulon properties and generate new hypotheses. The entire workflow for generating these knowledge bases is publicly available on GitHub (https://github.com/avsastry/modulome-workflow).

By systematically processing large-scale RNA-seq data, iModulonDB acts as a foundational resource for informing advanced experimental designs in metabolic engineering and broader synthetic biology applications. Thus, the transcriptomic datasets used in the present study are retrieved from this database when possible (for *E. coli* and *P. putida*). For the case of *Synechocystis sp. PCC 6803,* as no entry is available, we reproduced the transcriptomics dataset extraction and pre-processing steps described in the original paper^62^.

### iModulonDB pre-processing pipeline

For the case of *Synechocystis sp. PCC 6803,* no data was available on *i*ModulonDB. Therefore, to obtain transcriptomic data for this organism with comparable quality than the other 2 selected organisms, we executed the same pipeline used for iModulonDB construction.

The iModulonDB pre-processing pipeline systematically extracts and curates RNA-seq data to build a comprehensive transcriptional regulatory network (TRN) knowledge base. The pipeline commences by compiling sequence data and its correspondent metadata for all publicly available RNA-seq data from NCBI SRA database. After that, raw sequencing data is processed into log-transformed Transcripts Per Million (log-TPM) using a suite of bioinformatics tools including Trim Galore, FastQC, Bowtie, RSEQC, and featureCounts. Finally, a rigorous quality control phase filters out low-quality datasets based on metrics such as sequence quality and read mapping, complemented by manual curation of experimental metadata to ensure accuracy and consistency (see specific filters in Figure 12)^61^.

Importantly, batch effects, a common challenge when integrating data from diverse sources, are addressed by normalizing gene expression within each project against a defined reference condition, ensuring that observed data reflect biological rather than technical variation. However, for the contextualization of GEMs raw log-TPM values were used to avoid artifacts derived from normalization.

Applying this pipeline, 214 different transcriptomic datasets corresponding to RNA-seq sequencing data could be extracted. This amount is comparable to the *P. putida* dataset (321) and therefore we did not increase the search for more transcriptomic data using other methods. All code developed and used for this task can be seen inside the *Syneco_RNASeq_processing* folder of EMBER repository.

### Conditions selected for adaptability trade-off analysis

As mentioned before, to explore the trade-of between adaptability and growth in diverse conditions for addressing context-dependence, we use 3 different organisms. Namely, those organisms are *Escherichia coli K12*, *Pseudomonas putida KT2440* and *Synechocystis sp. PCC6803*).

Between those 3 organisms we have the representation of 3 different ecological niches, mammalian digestive system, soil and fresh water respectively. Those environments have unique dynamics and composition that would pose different pressures over the adaptability responses of organisms inhabiting them. Therefore, we should expect such responses to be diverse in both quantitative (as effort inverted) and quantitative (type of adaptation) among the selected organisms.

However, those habitat adaptations also make it difficult for the comparation of the trade-off between the organisms, as the same conditions could not be applied for all of them (i.e. autotrophy conditions are feasible for *Synechocystis* but not for *E. coli* or *P. putida*). In this way, we have tried to select relevant conditions for studying adaptations in carbon use, response to typical stresses of each organism or disruption on normal metabolic processes. All conditions tested and the criteria used are in the previous table.

### Gene classification

EMBER computes the association of a given reaction to biomass-related function (BAR) or to an adaptive response (NBR). However, to systematically characterize those sets we need to map this data to genes as, in contrast to reactions, they have consistent and high-quality functional annotation data. Classification of genes under these categories is made regarding the reaction presence in any of the mentioned categories.

At this point it is important to note that reaction presence is derived in GIMME from gene expression. As some reactions show complex GPR relationships, the mapping between gene expression and reaction presence is challenging. For example, in GIMME, the presence of reactions with *OR* GPRs is computed by assessing if the expression of the most expressed gene is above the imposed threshold. Although this method is effective for determining whether a reaction is active or not, it introduces a bias towards reaction inclusion by favoring those with complex *OR* GPRs involving several genes, compared with single-gene reactions. Consequently, it can happen that a single-gene reaction is determined to be inactive (and not included in the contextualized GEM) despite its gene being active according to expression data (from now on GIMME bias).

We detected this bias generation and acted to account for its impact. Considering it, GIMME will include all necessary reactions needed for meeting that the BOF is above the determined threshold. Hence, for the GIMME bias case, a reaction will only be excluded if the function of that reaction is not aligned with BOF. Therefore, for solving the introduced artifact, we classify genes that suffer GIMME bias as NBR.

For the rest of the genes, we will classify them according to their reaction presence in NBR or BAR. As discussed previously, we assessed reaction presence consistency across a threshold range from the 5th to the 20th percentile. In this way, reactions present in a set above 50% of the imposed percentiles are considered to be active. This is directly transferred to genes, with a given gene having a percentage of presence in BAR and NBR sets. Considering that, genes present in any reaction with a presence above 50% in the BAR set are considered of this class for a given condition. On the contrary, a gene whose associated reactions have a presence above 50% in the NBR set will be classified as such, conforming what we will call NBR genes or adaptative genes.

All this workflow was coded by the python function *get_bet_hedging_genes*, which is defined in the *Find_candidate_genes.ipynb* file within the *EMBER* repository. For an example of gene classification see Figure 13.

**Figure 13.**
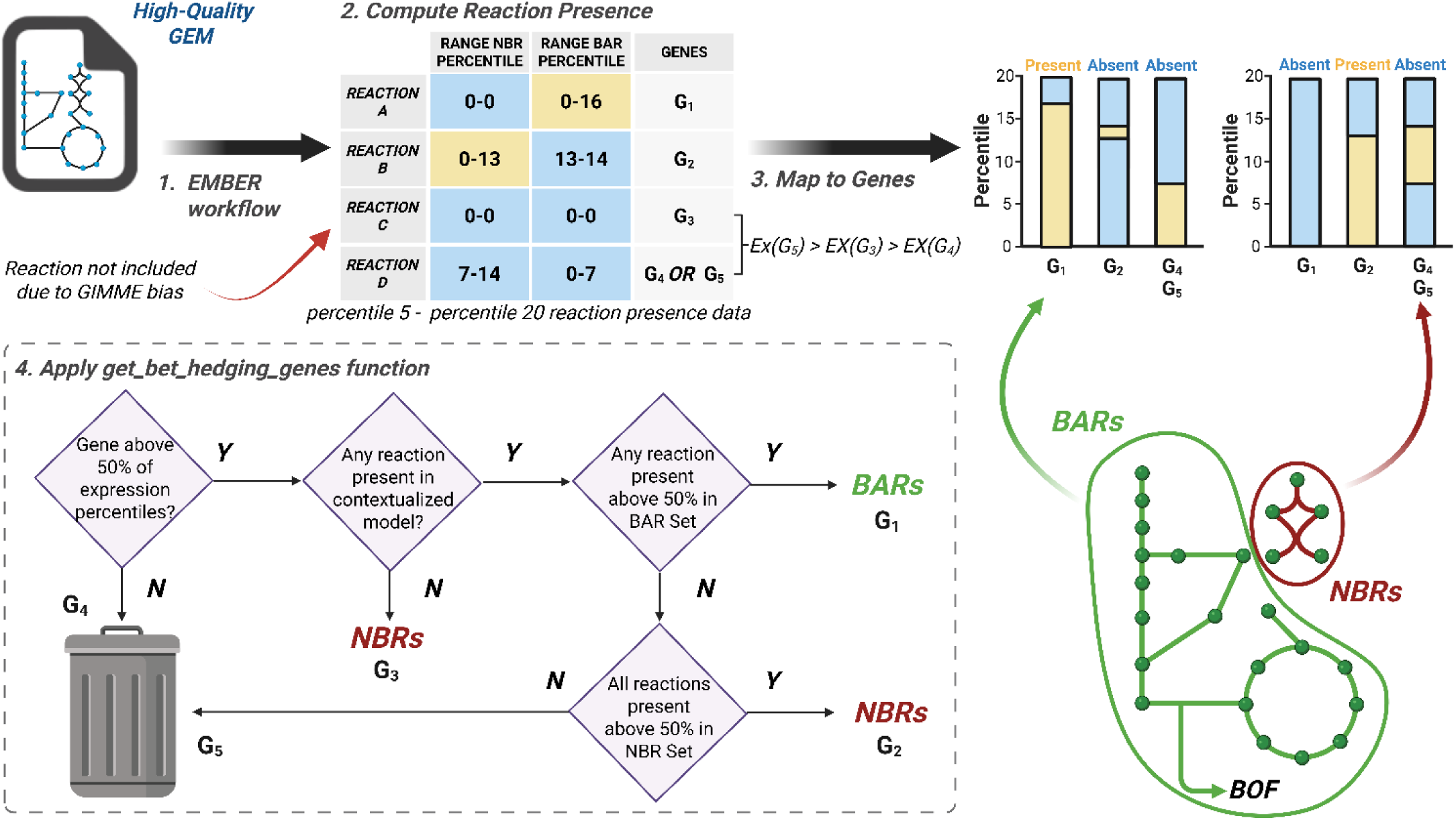
Schematic example of gene classification workflow used in this work. An idealized example of the classification of 5 genes according to the proposed workflow. In this example, the presence data for 4 reactions with different GPRs is computed through *EMBER* for a given condition, deriving active (yellow) or unactive reactions (blue) for each of reaction sets. In this step, GIMMIE bias favors the inclusion of reactions with complex GPRs due to their reaction expression mapping. This enables genes with higher expression values to end up with smaller presence values than others with less expression but included in a reaction with more complex GPR (exemplified with G3 and G4 respectively). After reaction presence it is computed is mapped directly to genes, determining its presence in each reaction set. Eventually, each gene is classified by using the function *get_bet_hedging_genes*, whose decision making is recapitulated in the presented flowchart. Note that despite G5 having an expression value enabling its inclusion up to the percentile 14, as it does not pass the 50% threshold for any of the reaction sets, it is discarded for consideration.

### Generation of homogeneous annotations with RAST

RAST server was developed as a fully automated service for annotating bacterial and archaeal genomes, aiming to rapidly produce high-quality assessments of gene functions and initial metabolic reconstructions. Its approach relies on a continually growing library of manually curated subsystems and protein families (FIGfams), which form the basis for its assertions of gene function and metabolic reconstruction^17,74^. For example, RAST annotates uploaded contigs in FASTA format, integrating the results into the SEED environment for comparative analysis.

This service can provide insights that fill gaps in required metabolic pathways, such as identifying a gene for menaquinone biosynthesis in *Mycobacterium tuberculosis*^75^. When compared with the KAAS (KEGG Automatic Annotation Server) service, RAST demonstrated comparable annotation quality, particularly in projecting existing SEED annotations. Discrepancies between RAST and manually curated SEED annotations were often reconcilable, with RAST missing only a small percentage of non-hypothetical genes^17^.

In the present work, the RAST server was used for generating a coherent and comparable annotation between the 3 selected species for performing a functional analysis of metabolic heterogeneity. This was performed by applying the default setup of RAST server (https://rast.nmpdr.org/) for the genome sequences in Table 3.

**Table 3.** Genome sequences used as input for RAST annotation.

| **NCBI Entry** | **Type** | **Organism** | **Size** |
| --- | --- | --- | --- |
| NC_000913.3 | Complete genome | *Escherichia coli str. K-12 substr. MG1655* | 4.64 Mb |
| NC_002947.4 | Complete genome | *Pseudomonas putida KT2440* | 6.18 Mb |
| NC_000911.1 | Complete genome | *Synechocystis sp. PCC 6803* | 3.57 Mb |

### Enrichment analysis

The rationale behind an enrichment analysis is to provide biological meaning to statistically derived groups by testing whether they are disproportionately associated with known functional or regulatory categories. When *EMBER* is applied to transcriptomic data, it produces sets of reactions varying together across conditions. While these groups (BARs and NBRs) capture coherent transcriptional signals, their biological significance is not immediately apparent. Enrichment analysis addresses this gap by comparing the computed groups to established annotation sets, such as transcription factor regulons, metabolic pathways, or Gene Ontology terms, to see if the overlap is greater than would be expected by random chance. The underlying assumption is that if a given group contains a statistically significant excess of biological entities from a known functional category, it likely represents the activity of a specific regulatory mechanism or biological process. By systematically performing this comparison across all computed groups, enrichment analysis transforms abstract mathematical components into interpretable biological modules. This enables us to link data-driven discoveries to existing knowledge and to generate new hypotheses about gene regulation, function, and network organization.

In the present work we use the implementation of gene set enrichment analysis (GSEA) developed in the pyModulon python package (https://pymodulon.readthedocs.io/). In PyModulon, enrichment analysis is used to determine whether the set of genes within an independently modulated gene set identified through Independent Component Analysis (ICA) is significantly associated with predefined biological categories such as transcription factor regulons, functional annotations (e.g., COG, GO terms), or user-defined gene sets. For each set–annotation pair, PyModulon applies a one-sided Fisher’s exact test to evaluate whether the observed overlap between the two sets is greater than expected by chance. This is done by constructing a contingency table based on the presence or absence of genes in each set. Because many tests are performed, the package controls for false positives by applying the Benjamini–Hochberg False Discovery Rate (FDR) correction, using a default significance threshold of q-value < 0.05. The results are returned as a dataset containing the gene set name, the annotation set name, raw p-value, FDR-adjusted q-value, number of overlapping genes, and the list of those genes^62^.

In our analysis we performed two different GSEAs. One with the reactions sets directly derived from EMBER and other for genes mapped from these reactions. In the approach concerning reactions, we use the subsystems of the GEMs as the functional categories. Instead, for the gene approach we used annotations derived from RAST as explained in the previous section. For both cases, two different thresholds were considered to decide if a set is enriched on a function or not. In this way, a set whose q-value is above 0.05 is considered not to be enriched. On the other hand, sets with q-value ≤ 0.05 and q-value ≤ 0.01 are considered as enriched and highly enriched sets respectively.

As RAST categories are hierarchical, in the GSEA concerning gene sets we performed a decomposition analysis of the functional categories which measures the specificity of the enrichment. For this, different levels of RAST functional categories were used as the target set in the analysis.

## ACKNOWLEDGEMENTS

This research was funded by the European Union’s Horizon 2020 research and innovation program under grant agreements 870294 (MIX-up), 101081782 (deCYPher) and 101112378 (PROMISEANG); Spanish Ministry of Science, Innovation and Universities (AEI/10.13039/501100011033) through research grant Rob3D (PID2022-139247OB-I00) and RobExplode (PID2019-108458RB-I00). AGB acknowledges funding from a predoctoral contract (FPI_PRE2020-092257).

## Notes

### Competing Interest Statement

The authors have declared no competing interest.

